# Non-cell-autonomous regulation of germline proteostasis by insulin/IGF-1 signaling via the intestinal peptide transporter PEPT-1

**DOI:** 10.1101/2024.02.22.581543

**Authors:** Tahir Muhammad, Stacey L. Edwards, Allison C. Morphis, Mary V. Johnson, Vitor De Oliveira, Tomasz Chamera, Siyan Liu, Ngoc Gia Tuong Nguyen, Jian Li

## Abstract

Gametogenesis involves active protein synthesis and heavily relies on proteostasis. How animals regulate germline proteostasis at the organismal level is poorly understood. Taking *C. elegans* as a model, we show that germline proteostasis requires coupled activities of HSF-1-dependent protein folding and insulin/IGF-1 signaling controlled protein synthesis. Depletion of HSF-1 from germ cells impairs chaperone gene expression, causing protein degradation and aggregation and, consequently, declines in fecundity and gamete quality. Reduced insulin/IGF-1 signaling confers germ cells’ resilience to limited protein folding capacity and proteotoxic stress by lowering ribosome biogenesis and the rate of translation. Interestingly, insulin/IGF-1 signaling promotes the expression of the evolutionarily conserved intestinal peptide transporter PEPT-1 via its downstream transcription factor FOXO/DAF-16, therefore allowing dietary proteins to be incorporated into an amino acid pool that fuels protein synthesis in the germline. We propose that this non-cell-autonomous pathway plays a critical role in regulating proteostasis in gametogenesis.

**Teaser:** Insulin/IGF-1 signaling regulates proteostasis in gametogenesis via the control of dietary protein absorption.

## Introduction

Gametogenesis, in which primordial germ cells develop into gametes (sperm or oocytes) through mitotic proliferation, meiotic differentiation, and gamete maturation, is essential for sexually reproducing animals to transmit genetic information over generations while introducing genetic diversity (*1*). During gametogenesis, germ cells experience periods of very active protein synthesis to make cell-cycle-specific proteomes through mitosis and meiosis and to prepare maternal proteins for embryogenesis (*2, 3*). These newly synthesized proteins must be folded into proper conformations assisted by molecular chaperones to be functional (*4*). How animals coordinate protein translation and folding during gametogenesis to ensure sufficient folding capacity is poorly understood.

Heat shock factor 1 (HSF1), a key transcription regulator of cellular responses to proteotoxic stress (*5, 6*), has evolutionarily conserved roles in gametogenesis in both vertebrates and invertebrates (*7–9*). It activates the transcription of selective molecular chaperones and co-chaperones in germ cells under physiological and stress conditions (*9–12*). Reports from *C. elegans* and mice also suggest that HSF1 binds to the promoters and regulates the expression of genes with other essential functions in reproduction including meiosis (*9, 13*). It is yet to be determined whether the primary role of HSF1 in germline development is to maintain proteostasis.

Our previous study has shown that the requirement for HSF-1, the only HSF in *C. elegans,* in germline development is dictated by the activity of insulin/IGF-1 signaling (IIS) (*9*). While in the wild-type animals, loss of HSF-1 from the germline through larval development results in sterility, reduced IIS could partially restore fecundity in the absence of HSF-1 in germ cells. This finding implies that reducing IIS may rewire the proteostasis network and render germ cells less dependent on HSF-1-mediated chaperone expression. IIS is a highly conserved nutrient-sensing pathway that promotes gametogenesis both cell-autonomously in the germline and non-cell-autonomously from somatic tissues (*3, 14*). In addition, IIS activity needs to be fine-tuned for gamete quality as hyperactivation of IIS has detrimental effects on oogenesis while reduced IIS preserves functional oocytes during maternal aging in mice and *C. elegans* (*15, 16*). On the other hand, IIS impacts proteostasis and responses to proteotoxic stress through transcriptional and post-transcriptional control of many players in protein synthesis, turnover, and quality control (*17–21*). However, it is not clear why the activities of HSF-1 and IIS need to be coupled for germline development and how IIS regulates germline proteostasis at the organismal level.

In this study, we found that HSF-1 is important for germline proteostasis by enhancing protein folding capacity in *C. elegans* gametogenesis at ambient temperature. Reduced IIS grants germ cells resilience against proteotoxic challenges associated with loss of HSF-1 and stress through lowering ribosomal biogenesis and translation. Interestingly, IIS promotes protein synthesis in the germline non-cell-autonomously via transcriptional activation of the intestinal peptide transporter, PEPT-1. Our findings suggest that IIS-mediated dietary protein absorption and germline protein synthesis must work in concert with HSF-1-dependent protein folding to maintain proteostasis in gametogenesis.

## Results

### HSF-1 is important for germline proteostasis at ambient temperature

HSF-1 is well-known for its roles in proteotoxic stress responses, such as against heat shock, but it also regulates gene expression in physiological conditions during development, reproduction, and aging (*9, 22, 23*). In a previous study, we identified the HSF-1 transcriptional program in *C. elegans* germline by combining tissue-specific HSF-1 depletion using the auxin-inducible degron (AID) system with ChIP-seq and RNA-seq analyses at ambient temperature (*9*). Among those genes that had HSF-1 binding at the promoters and decreased expression upon HSF-1 depletion in the germline of young adults are proteostatic and reproduction-related genes. Kinetic analysis indicated that a group of chaperone and co-chaperone genes were the most sensitive to HSF-1 depletion by showing the quickest decline of mRNA levels (*9*), implicating that they are likely the primary target genes of HSF-1 in gametogenesis. Interestingly, following the decrease of chaperone mRNA expression at 8 h of HSF-1 depletion from the germline, genes in the ubiquitin-proteasome system (UPS) were induced at the 16 h time point (Fig. 1A). This result implicates that the UPS is upregulated as a stress response to remove misfolded proteins caused by loss of HSF-1-dependent chaperone expression and decrease in protein folding capacity.

**Figure 1.**
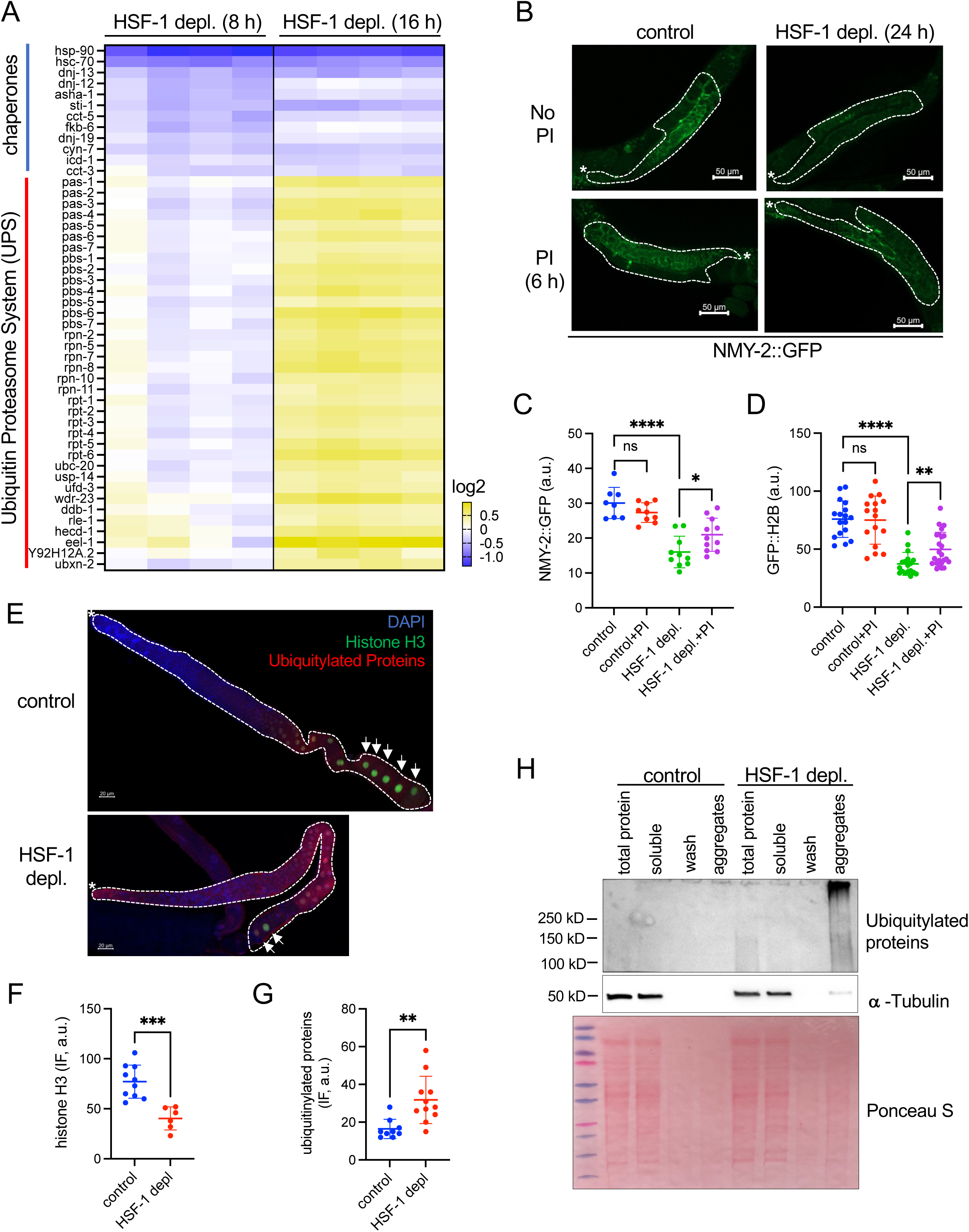
Loss of HSF-1 from the germline impairs the expression of chaperone genes, causing protein degradation and aggregation. (A) Heatmap of mRNA fold changes at HSF-1-dependent chaperone and co-chaperone genes as well as components of the ubiquitin-proteasome system (UPS) upon depletion of HSF-1 (HSF-1 depl.) from the germline of young adults using auxin-inducible degradation (AID) for 8 h and 16 h. Differentially expressed genes (FDR: 0.05) at either time point in both functional groups are included. The promoters of chaperone and co-chaperone genes shown in the heatmap are bound by HSF-1 in germ cells as determined by ChIP-seq. (B&C) Representative images (B) and quantification of protein levels of NMY-2::GFP transgene (C) in the germline upon HSF-1 depletion. HSF-1 was depleted from the germline of young adults using AID for 24 h in the presence or absence of the proteasome inhibitor bortezomib (PI) for the last 6 h of HSF-1 depletion. The dashed lines outline the gonads. The white asterisks (*) indicate the distal end of gonads where progenitor cells are located. (D) Quantification of protein levels of GFP::H2B transgene in fully grown oocytes upon germline-specific depletion of HSF-1. Experiments were done as in B&C. (E&F) Representative images (E) and quantification (F) of endogenous histone H3 and ubiquitylated proteins (not free ubiquitin) by immunofluorescence (IF) upon germline-specific depletion of HSF-1 in young adults for 24 h. The dashed lines outline the gonads with the white asterisks (*) marking the distal end. The arrows indicate the fully grown oocytes, where the levels of histone H3 are quantified. (H) Western blot of protein aggregates upon germline-specific depletion of HSF-1 in young adults for 24 h. Antibodies against ubiquitin and α-tubulin were used. The ‘aggregates’ fraction was loaded as 10-fold of the other fractions. In all the dot plots, the mean and standard deviation are plotted. Student’s t-test. *p<0.05; **p<0.01; ***p<0.001; ****p<0.0001; ns: p>=0.05.

To test this hypothesis, we took the GFP fusion of NMY-2 (non-muscle myosin) that resides in the cytosol of the germline (Fig. 1B) and histone H2B that accumulates in the nuclei of oocytes (Fig. S1A) as examples to test protein stability upon HSF-1 depletion. Loss of HSF-1 from the germline for 24 h led to a significant decrease of both NMY-2 and H2B, which were partially reversed by inhibition of proteasome for 6 h before imaging analysis (Fig. 1B-D & Fig.S1B). These results suggest that HSF-1-mediated chaperone expression is essential for the stability of both cytosolic and nuclear proteins in germ cells. Notably, the short proteasome inhibition was not sufficient to restore HSF-1 protein levels from the depletion by AID (Fig. S1B). Therefore, the partial recovery of GFP fusion proteins was not due to re-gaining HSF-1 activities in the germline but rather through preventing the degradation of NMY-2 and H2B fusion proteins by the UPS. The endogenous histone H3 showed a similar decline in protein levels upon HSF-1 depletion as measured by immunofluorescence (IF) (Fig. 1E&F). In addition, we monitored the change of ubiquitylated proteins in the germline upon loss of HSF-1. Low levels of ubiquitylated proteins were visible in the presence of HSF-1 at the diplotene stage and in oocytes, consistent with the reported functions of protein ubiquitylation in meiosis (*24, 25*). Upon HSF-1 depletion, levels of ubiquitylated proteins significantly elevated through the germline (Fig. 1E&G) suggesting the increase of misfolded proteins ubiquitinylated for clearance. Finally, we found that high molecular weight, ubiquitylated proteins increased in the detergent-insoluble aggregates upon HSF-1 depletion from the germline, supporting them as the misfolded species and aggregation-prone (Fig. 1H). Interestingly, we also found that a small fraction of α-tubulin was in the insoluble fraction upon HSF-1 depletion, indicating the gain-of-function toxicity of misfolded proteins that trapped other proteins into aggregates and further imbalanced the proteome. The accumulation of ubiquitylated proteins in aggregates also suggests the protein degradation system may be overwhelmed by the continuous influx of misfolded proteins due to insufficient folding capacity. Supporting this notion, the levels of NMY-2::GFP were recovered after a prolonged 48 h depletion of HSF-1 (Fig. S1C-E) with the appearance of GFP in puncta-like structures (Fig. S1D, lower panel).

Collectively, our data demonstrate that protein misfolding, degradation, and aggregation increase following the decline of HSF-1-dependent chaperone expression, indicating that HSF-1 is important for providing protein folding capacity in germ cells.

### Insufficient protein folding capacity compromises fecundity and gamete quality

We then tested the impacts of insufficient protein folding capacity on gametogenesis. As soon as 16 h of HSF-1 depletion from young adult germline, we observed a significantly increased number of nuclei in the mitotic zone where germline stem cells (GSCs) are located (Fig. 2A&B), but a smaller percentage of them were in the S-phase (EdU positive, Fig. 2A&C). These results suggest that loss of HSF-1 impairs GSC proliferation and its progression into meiotic prophase I. The defects that we observed in gametogenesis very likely resulted from the insufficient protein folding associated with the early decline of HSF-1-dependent chaperone and co-chaperone genes, as at this time point, the other genes that have HSF1 binding at the promoters in the germline just began changing their mRNA expression. Consistent with the cellular defects, we found the fecundity of self-reproducing hermaphrodites declined dramatically starting from the second day of HSF-1 depletion (Fig. 2D). We conclude that it was due to the impairment of ongoing oogenesis in young adults rather than the potential impacts on the quality of sperm already made at the L4 larval stage because a similar decline of fecundity was observed upon HSF-1 depletion when we mated the hermaphrodites with wild-type males (Fig. 2E). Loss of HSF-1 from the germline and the consequent decline in protein folding also compromised the quality of oocytes as embryo lethality increased by more than 10-fold in both self progenies and mated progenies starting from the second day of HSF-1 depletion (Fig. 2F).

**Figure 2.**
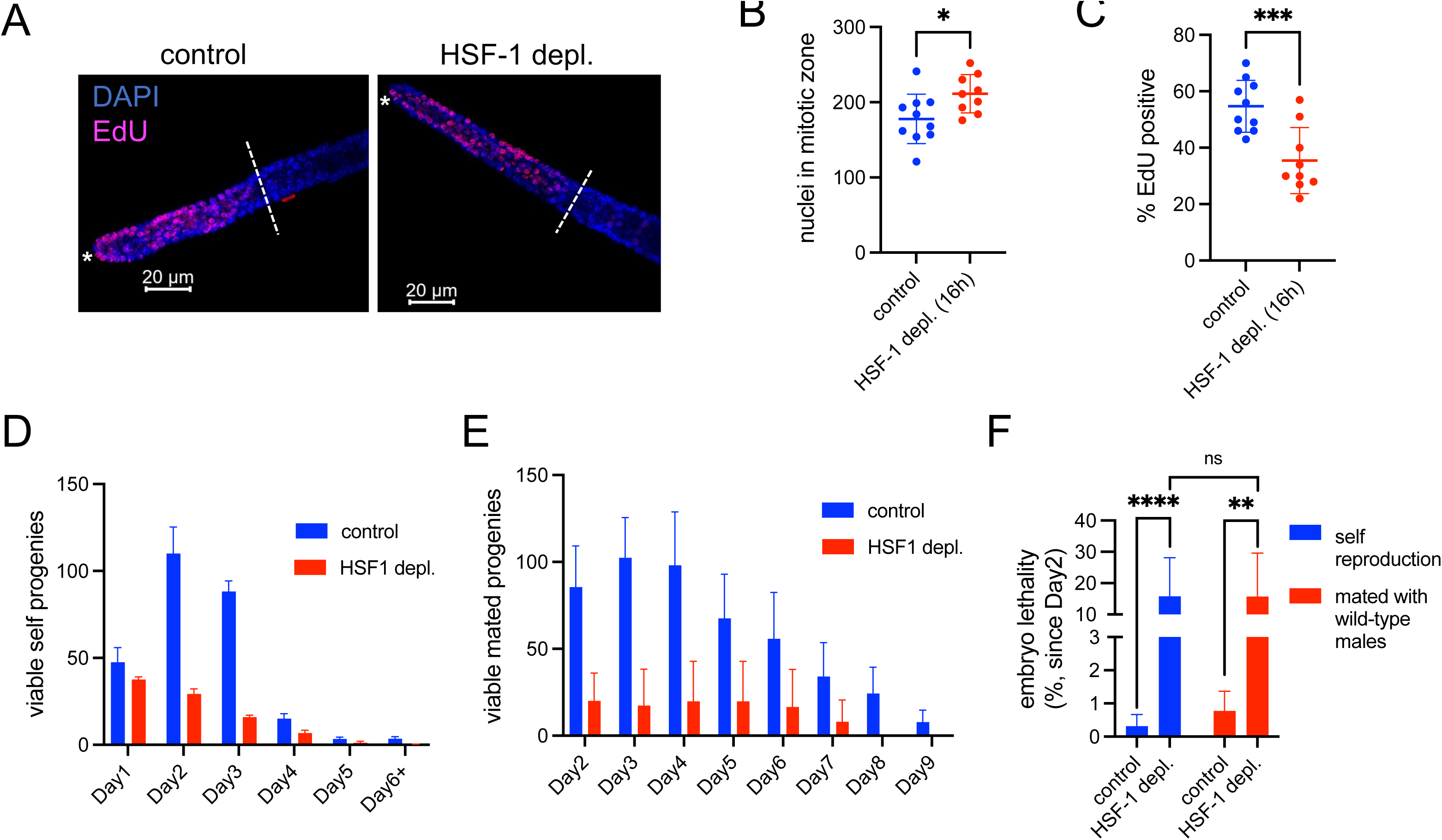
HSF-1 is required in the adult germline for fecundity and oocyte quality. (A) Representative images of gonads with EdU labeling of nuclei in the S-phase and DAPI staining of all germline nuclei upon depletion of HSF-1 from the germline of young adults for 16 h. Progenitor cell proliferation occurs in the mitotic zone between the distal end of gonads marked by the white asterisks (*) and the dashed lines that indicate where the nuclei start transitioning into meiotic pre-phase I. (B&C) Quantification of the total number of mitotic nuclei (B) and percent in the S-phase (C) upon depletion of HSF-1 from the germline of young adults for 16 h. Mean and standard deviation are plotted. Student’s t-test. *p<0.05; ***p<0.001. (D) Fecundity as measured by self-progenies when HSF-1 was depleted from the germline of hermaphrodites starting from Day 1 of adulthood. Mean and SEM are plotted (three experiments, N>=12 for each experiment). (E) Fecundity as measured by mated progenies when HSF-1 was depleted from the germline of hermaphrodites and mated with N2 males on Day 1 of adulthood. Mean and standard deviation are plotted (control: N=12; HSF-1 depletion: N=11). (F) Embryo lethality caused by depletion of germline HSF-1. HSF-1 was depleted from the germline of hermaphrodites starting from Day 1 of adulthood, and embryo lethality in self-progenies and mated progenies (with N2 males) was measured on Day 2 and after. Mean and standard deviation are plotted (self-reproduction: N=15; mating experiments: N>=10). Student’s t-test. **p<0.01; ****p<0.0001; ns: p>=0.05.

### Reduced insulin/IGF-1 signaling confers germ cells’ resilience against limited protein folding capacity

Despite HSF-1’s role in germ cell proteostasis, our previous study showed that reduced insulin/IGF-1 signaling (IIS) could alleviate the requirement for HSF-1 in germline development. When HSF-1 was depleted from the germline through larval stages, the wild-type animals were utterly sterile, while the reduction-of-function mutant of insulin/IGF-1 receptor, daf-2(e1370), partially restored the reproduction (*9*). Similar results were observed when we depleted HSF-1 from the germline at the young-adult stage after hermaphrodites switched from spermatogenesis to oogenesis (Fig. 3A&B). Reduced IIS as in the *daf-2(e1370)* animals or *daf-2(e1370)* animals further treated with *daf-2* RNAi decreased the brood size in the presence of HSF-1 (Fig. 3A), consistent with the role of IIS in promoting gametogenesis (*26, 27*). However, reduced IIS rendered fecundity less dependent on HSF-1 (Fig. 3A) and dramatically decreased embryo lethality associated with HSF-1 depletion (Fig. 3B). These results implicate that reduced IIS could improve proteostasis in germ cells without HSF-1.

**Figure 3.**
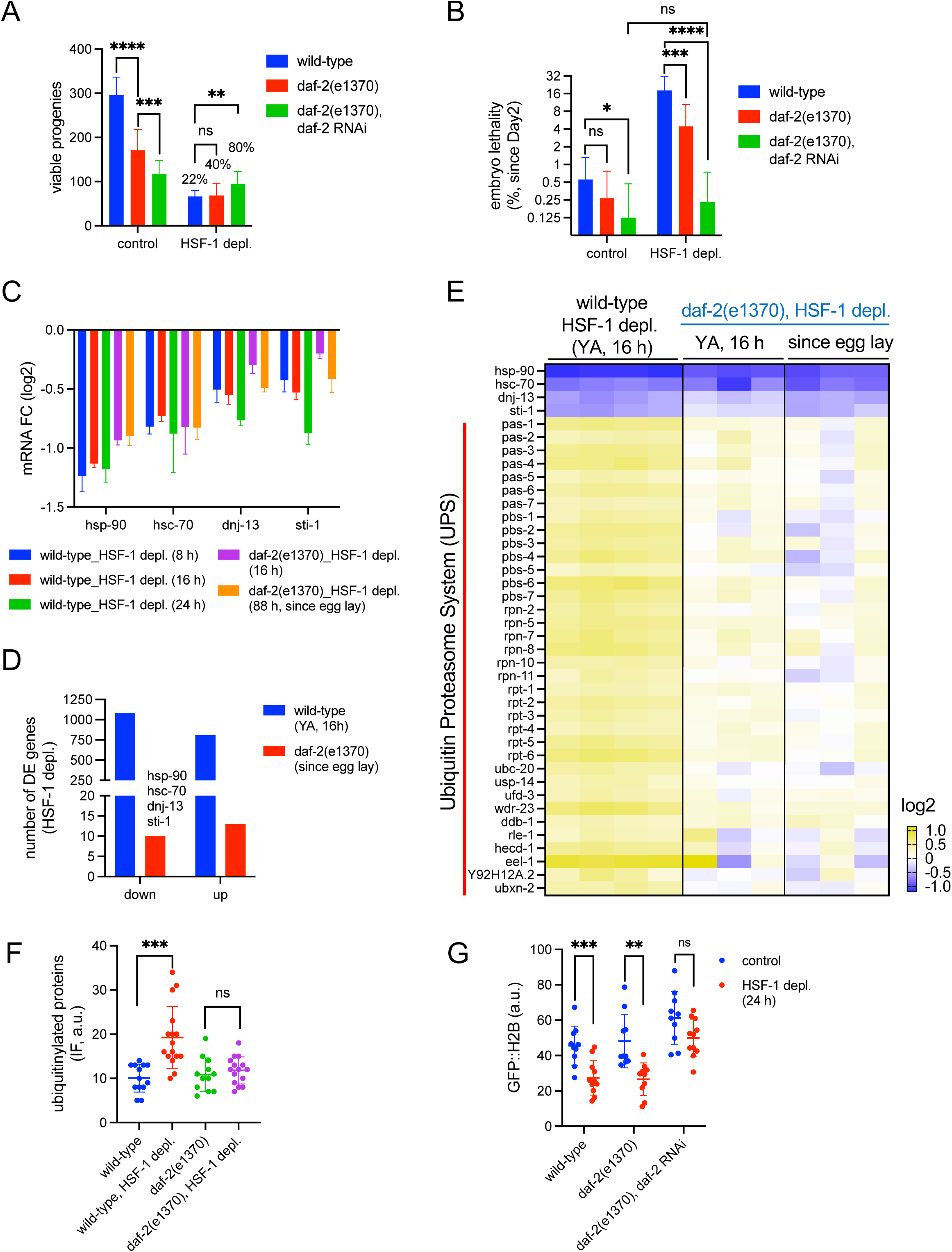
Reduced Insulin/IGF-1 signaling (IIS) confers resilience against limited protein folding capacity in gametogenesis. (A&B) Brood size (A) and embryo lethality (B) of animals with reduced IIS upon depletion of HSF-1 from the germline starting from Day 1 of adulthood. The wild-type and *daf-2(e1370)* animals were treated with daf-2 RNAi or control RNAi (L4440) starting at egg lay. Embryo lethality was measured on Day 2 and after. Mean and standard deviation are plotted (N>=15). The percentage of brood size (HSF-1 depletion compared with the control) is labeled. Student’s t-test. *p<0.05, **p<0.01; ***p<0.001; ****p<0.0001; ns: p>=0.05. (C) Histograms showing mRNA fold changes (FC) of selective chaperone and co-chaperone genes upon HSF-1 depletion from the germline as measured by RNA-seq analysis in the wild-type and *daf-2(e1370)* animals. The four chaperone and co-chaperone genes showed significant changes in mRNA levels (FDR: 0.05) upon HSF-1 depletion in wild-type and *daf-2(e1370)* animals. In the wild-type animals, HSF-1 was depleted from the young adults for 8 h, 16 h, or 24 h. In the *daf-2(e1370)* animals, HSF-1 depletion was done in the young adults for 16 h or started from egg lay for 88 h till the animals grew to gravid adults. Mean and standard deviation are plotted (wild-type 8 h & 16 h: N=4; wild-type 24 h & *daf-2(e1370)* experiments: N=3). (D) Histograms showing the number of differentially expressed (DE) genes (FDR: 0.05) caused by HSF-1 depletion from the germline in the wild-type animals at young-adult (YA) stage for 16 h and from the *daf-2(e1370)* animals starting from egg lay. (E) Heatmap of mRNA fold changes at selective chaperone and co-chaperone genes as well as components of the ubiquitin-proteasome system (UPS) upon depletion of HSF-1 from the germline of the wild-type and *daf-2(e1370)* animals. (F) Quantification of ubiquitylated proteins in the gonads by immunofluorescence (IF) upon germline-specific depletion of HSF-1 for 24 h from young adults of the wild-type and *daf-2(e1370)* animals. Mean and standard deviation are plotted. Student’s t-test. ***p<0.001; ns: p>=0.05. (G) Quantification of protein levels of GFP::H2B transgene in fully grown oocytes upon germline-specific depletion of HSF-1 for 24 h from young adults. The wild-type and *daf-2(e1370)* animals were treated with daf-2 RNAi or control RNAi (L4440) starting from egg lay. Mean and standard deviation are plotted. Student’s t-test. **p<0.01; ***p<0.001; ns: p>=0.05.

To understand why IIS dictates the requirement for HSF-1 in germ cells, we first tested if reduced IIS restores chaperone expression in the absence of HSF-1 by RNA-seq analysis. The genes encoding two essential ATP-dependent chaperones, *hsp-90* and *hsc-70* (also called *hsp-1*), and two critical co-chaperones, *dnj-13* and *sti-1*, were still significantly down-regulated when HSF-1 was depleted from the germline of *daf-2(e1370)* animals (Fig. 3C&D). Significantly, upon chronic depletion of HSF-1 starting from egg lay in the *daf-2(e1370)* animals, the decrease in mRNA levels of those chaperone and co-chaperone genes was comparable with the 16 h of HSF-1 depletion in the wild-type animals (Fig. 3C) that was sufficient to cause gametogenesis defects (Fig. 2A-C) and expression changes at ∼2000 genes in the wild-type animals (Fig. 3D).

We then tested if reduced IIS mounts a more robust stress response to loss of HSF-1 from the germline. Reduced IIS enhances stress response through downstream transcription factors, such as FOXO/DAF-16 and NRF/SKN-1 (*19, 28–30*). Our previous study demonstrates that the tolerance to germline HSF-1 depletion by reduced IIS is independent of SKN-1 and only requires DAF-16 non-cell-autonomously from the soma (*9*), suggesting that the stress responses mediated by SKN-1 and DAF-16 unlikely play essential roles in the germline to resolve proteotoxicity from loss of HSF-1. Our RNA-seq analysis showed that the UPS components were not induced either upon acute or chronic depletion of HSF-1 from the germline of *daf-2(e1370)* animals (Fig. 3E). Furthermore, loss of HSF-1 from germ cells only altered a handful of genes in addition to the four chaperone and co-chaperone genes mentioned above in the *daf-2(e1370)* animals (Fig. 3D). These results implicate the intrinsic property of germline proteome in the *daf-2(e1370)* animals rather than stress response underlies its resilience against low protein folding capacity.

Finally, we tested if reduced IIS enhances proteostasis in germ cells without HSF-1. Consistent with the expression data on the UPS components, ubiquitylated proteins did not significantly increase in the germline of *daf-2(e1370)* animals upon HSF-1 depletion as in the wild-type animals, implicating less misfolded proteins resulted from HSF-1 depletion when IIS was reduced (Fig. 3F, S2A). Although the *daf-2(e1370)* mutation was not sufficient to maintain histone protein levels in the oocytes without HSF-1 (Fig. 3G, S2A-C), further reducing IIS by *daf-2* RNAi restored the levels of GFP::H2B (Fig. 3G, S2C). This is consistent with the observation that HSF-1 depletion had minimal impacts on fecundity and embryo lethality in the *daf-2(e1370)* animals treated with *daf-2* RNAi (Fig. 3A&B). Collectively, our results indicate that reduced IIS confers germ cells’ resilience against compromised protein folding capacity associated with loss of HSF-1.

### Insulin/IGF-1 signaling (IIS) activates ribosome biogenesis and translation in germ cells

Next, we attempted to understand how reduced IIS remodels the germline proteostasis network by transcriptomic analysis in isolated germline nuclei from the *daf-2(e1370)* and the wild-type animals (Fig. S3). Genes involved in protein synthesis are among the most enriched functional groups that decreased expression (Fig. 4A), which include those encoding protein components of both ribosomal subunits, translation initiation and elongation factors, and one mitochondrial ribosomal protein gene (Fig. 4B). This result indicates that the translation machinery is coordinately down-regulated when IIS is low. Consistent with transcriptomic data, the protein levels of endogenously tagged RPS-6::mCherry significantly decreased in the *daf-2(e1370)* germline compared to that in the wild-type control, suggesting that IIS activates ribosome biogenesis in germ cells (Fig. 4C&D). Finally, we found the translation levels in the germline were dependent on IIS activity. Translation, as measured by O-propargyl-puromycin (OPP) incorporation, was significantly decreased in the germline of *daf-2(e1370)* animals, and further reduced upon *daf-2* RNAi (Fig. 4E&F). On the contrary, *daf-16* RNAi in the *daf-2(e1370)* animals partially restored translation (Fig. 4E&F). As DAF-16 is an essential regulator of GSC proliferation downstream of IIS (*26, 31–33*), our results strongly suggest that activation of protein synthesis underlies the role of IIS in promoting gametogenesis.

**Figure 4.**
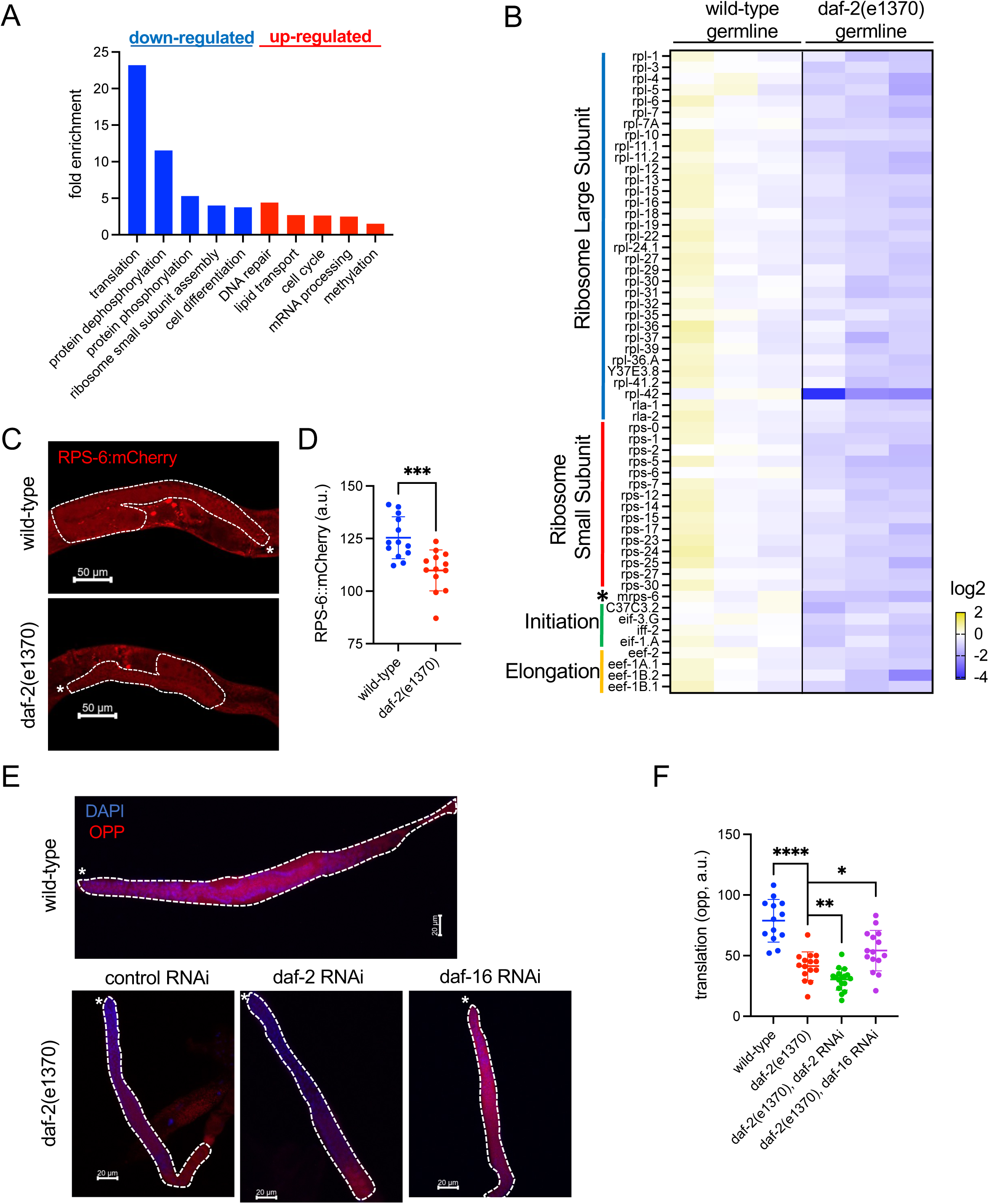
Insulin/IGF-1 signaling (IIS) activates ribosome biogenesis and translation in germ cells. (A) Gene Ontology (GO) analyses of differentially expressed (DE) genes (FDR: 0.05) when comparing the germline of the *daf-2(e1370)* animals to that in the wild-type animals. RNA-seq analysis was performed in isolated germline nuclei from young adults of the wild-type and *daf-2(e1370)* animals (N=3). The top 5 GO terms based on enrichment scores are shown for downregulated genes (blue bars) and upregulated genes (red bars) in the *daf-2(e1370)* germline. (B) Heatmap of the mRNA levels of translation-related genes in the wild-type and *daf-2(e1370)* germline. The fold change against the average of mRNA in the wild-type is shown. Genes that function in translation based on GO analysis and show significant changes in the *daf-2(e1370)* germline (FDR: 0.05) are included. The mitochondrial ribosomal protein gene, mrps-6, is labeled by *. (C&D) Representative images (C) and quantification (D) of protein levels of the endogenously tagged RPS-6::mCherry in the gonad of wild-type and *daf-2(e1370)* animals. The dashed lines outline the gonads, and the white asterisks (*) indicate the distal end of the gonads. Mean and standard deviation are plotted. Student’s t-test. ***p<0.001. (E&F) Representative images (E) and quantification (F) of protein translation in the germline of animals with altered IIS. Newly synthesized proteins were measured by O-propargyl-puromycin (OPP) incorporation. The wild-type and *daf-2(e1370)* animals were treated with daf-2 RNAi, daf-16 RNAi, or control RNAi starting from egg lay. Mean and standard deviation are plotted for quantification. Student’s t-test. *p<0.05, **p<0.01; ****p<0.0001.

### Tuning down protein translation enhances germline proteostasis against the challenges of protein misfolding

On the other hand, does a low rate of protein synthesis associated with reduced IIS render germline tolerant to limited protein folding capacity as upon HSF-1 depletion? To answer this question, we took *rsks-1(ok1255)*, the loss-of-function mutant of *C. elegans* ortholog of human p70 ribosomal S6 Kinase, as a model, which is known to reduce the rate of translation (*34*). The *rsks-1(ok1255)* animals limited the accumulation of protein misfolding as shown by the levels of ubiquitylated proteins in the germline (Fig. 5A, S4A), and partially stabilized the histone H3 in the oocytes upon HSF-1 depletion (Fig. S4A&B). Consistent with the measurements of proteostasis, the *rsks-1(ok1255)* animals displayed almost no additional defects in fecundity and gamete quality when HSF-1 was absent from the germline (Fig. 5B&C). These results support that a low translation rate underlies the enhanced germline proteostasis by reduced IIS.

**Figure 5.**
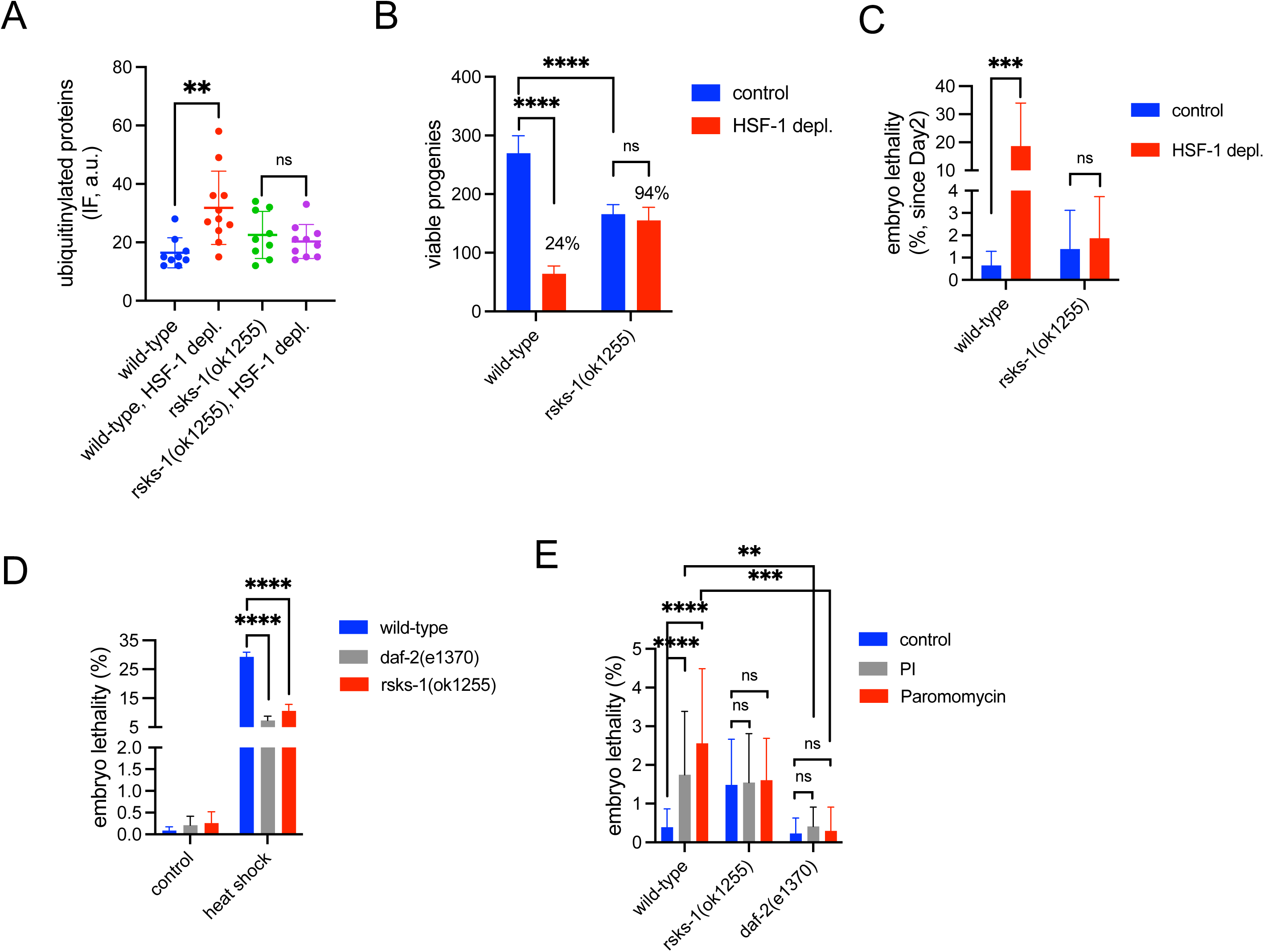
Reduced translation rate underlies robust germline proteostasis and resilience against proteotoxic stress mediated by low insulin/IGF-1 signaling (IIS). (A) Quantification of ubiquitylated proteins in the gonads by immunofluorescence (IF) upon germline-specific depletion of HSF-1 for 24 h from young adults of the wild-type and *rsks-1(ok1255)* animals. Mean and standard deviation are plotted. Student’s t-test. **p<0.01; ns: p>=0.05. (B&C) Brood size (B) and embryo lethality (C) of the *rsks-1(ok1255)* animals upon depletion of HSF-1 from the germline starting from Day 1 of adulthood. Embryo lethality was measured on Day 2 and after. Mean and standard deviation are plotted (wild-type control, N=10; wild-type HSF-1 depl., N=20; *rsks-1(ok1255)* control, N= 24; *rsks-1(ok1255)* HSF-1 depl., N=18). Student’s t-test. ***p<0.001; ****p<0.0001; ns: p>=0.05. (D) Embryo lethality caused by acute heat stress in the wild-type, *daf-2(e1370)* and *rsks-1(ok1255)* animals. Young gravid adults were heat-shocked briefly at 34 °C for 15 min, and embryo lethality was measured in eggs laid within 4 h of the recovery at 20 °C. Experiments were done in 8 biological replicates, each containing 5 animals. Mean and SEM are plotted. Student’s t-test. ****p<0.0001; ns: p>=0.05. (E) Embryo lethality caused by the proteasome inhibitor bortezomib (PI) and paromomycin that induces translational misreading in the wild-type, *daf-2(e1370)* and *rsks-1(ok1255)* animals. Animals were treated with either 10 μM of bortezomib or 1 mM of paromomycin starting at the young adult stage. Embryo lethality through the reproductive period was measured. Mean and standard deviation are plotted (N>=15). Student’s t-test. **p<0.01; ***p<0.001; ****p<0.0001; ns: p>=0.05.

Since our data suggest the primary role of HSF-1 in gametogenesis is to provide sufficient protein folding capacity, we further tested if reduced IIS and low translation rate could grant germ cells resilience against proteotoxic stresses that challenge folding. We first exposed the animals to acute heat stress, which is known to induce protein misfolding. A brief heat shock at 34 °C for 15 minutes was sufficient to lead to ∼30% embryo lethality from eggs laid within 4 hours of the stress in the wild-type animals. The *daf-2(e1370)* and *rsks-1(ok1255)* animals, however, reduced embryo lethality after heat shock by 75% and 64%, respectively (Fig. 5D). We then treated the animals with bortezomib, which inhibits proteasome and hinders the removal of misfolded proteins, and paromomycin, which induces translation misreading, during the entire reproductive period. These two stressors significantly increased embryo lethality in the wild-type animals but failed to do so in either *daf-2(e1370)* or *rsks-1(ok1255)* animals (Fig. 5E).

Collectively, our data suggest that tuning down translation by reduced IIS provides germline resilience against the challenges of protein misfolding either due to insufficient chaperone expression or acute and chronic proteotoxic stresses. Conversely, IIS-stimulated protein synthesis requires HSF-1-dependent protein folding capacity to maintain germline proteostasis.

### PEPT-1 functions downstream from insulin/IGF-1 signaling (IIS) and FOXO/DAF-16 to determine the requirement for HSF-1 in germline development

Our previous study showed that the requirement for HSF-1 in germline development is dictated by DAF-16 activity in the soma (*9*). Since DAF-16 is involved in the regulation of germline protein synthesis by IIS (Fig. 4E&F), which requires coupled activities of HSF-1 for chaperone expression, we hypothesize that DAF-16 through its transcriptional target genes in somatic tissues non-cell-autonomously impacts protein synthesis and proteostasis in the germline. To identify in which tissue DAF-16 mediates this non-cell-autonomous regulation, we performed tissue-specific RNAi against DAF-16 in the *daf-2(e1370)* animals and tested the sensitivity of germline development to HSF-1 depletion starting at egg lay (Fig. 6A). As we previously reported, the *daf-2(e1370)* animals were able to reproduce in the absence of germline HSF-1 through larval development, while systemic RNAi against *daf-16* led to sterility. The *daf-16* RNAi in the muscle or intestine significantly decreased the brood size of the *daf-2(e1370)* animals upon HSF-1 depletion but did not cause complete sterility, suggesting that DAF-16 functions from both tissues to regulate germline proteostasis. By contrast, *daf-16* RNAi in the hypodermis or germline did not significantly alter the brood size of the *daf-2(e1370)* animals upon HSF-1 depletion.

**Figure 6.**
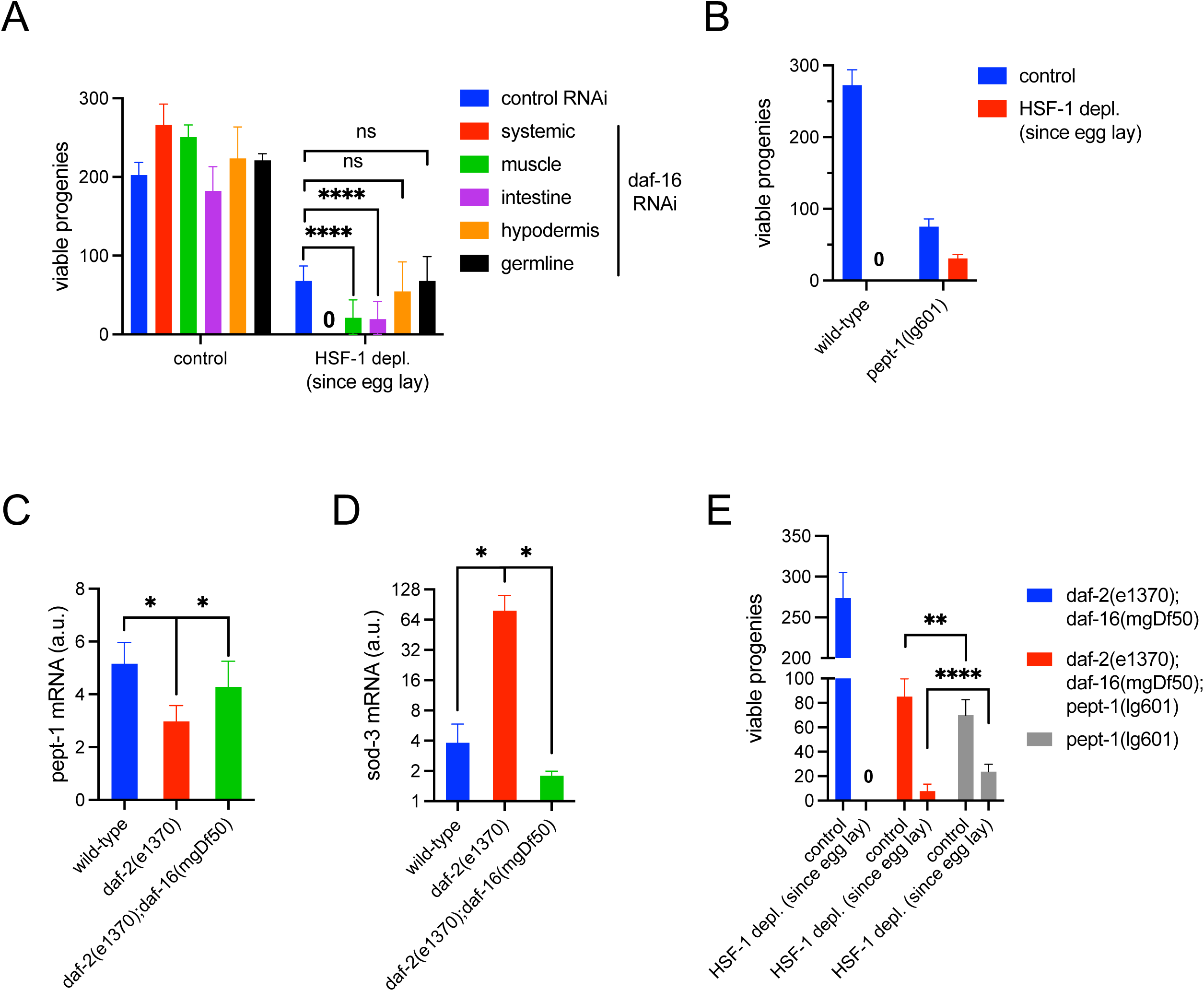
Intestinal peptide transporter PEPT-1 functions downstream from FOXO/DAF-16 to dictate the requirement for HSF-1 in germline development. (A) Brood size of the *daf-2(e1370)* animals treated with *daf-16* RNAi upon depletion of HSF-1 from the germline starting from egg lay. The *daf-2(e1370)* animals were treated with control RNAi (L4440) or *daf-16* RNAi either in whole animals (systemic) or in specific tissue starting from egg lay. Mean and standard deviation are plotted (N>=14). Student’s t-test. ****p<0.0001; ns: p>=0.05. (B) Brood size of the loss-of-function mutant *pept-1(lg601)* upon depletion of HSF-1 from the germline starting from egg lay. Mean and standard deviation are plotted (N>=14). (C&D) The RT-qPCR analysis of *pept-1* (C) and *sod-3* (D) in the wild-type, *daf-2(e1370)* and *daf-2(e1370);daf-16(mgDf50)* animals at late L4 stage. Mean and standard deviation of biological triplicates are plotted. Student’s t-test. *p<0.05. (E) Brood size of the *pept-1(lg601), daf-2(e1370);daf-16(mgDf50)* and *daf-2(e1370);daf-16(mgDf50); pept-1(lg601)* animals upon depletion of HSF-1 from the germline starting from egg lay. Mean and standard deviation are plotted (N>=14). Student’s t-test. **p<0.01; ****p<0.0001.

Based on transcriptomic data and DNA motif analysis, the DAF-16-responsive genes in the IIS pathway have been grouped into two classes: DAF-16 directly activates Class I genes through the DAF-16-binding element (DBE) and indirectly represses Class II genes that contain the DAF-16-associated element (DAE) (*35*). We decided to focus on a top-ranked Class II gene, *pept-1* in our study for the following reasons: first, it encodes an oligopeptide transporter specifically expressed in the intestine, one primary tissue from which DAF-16 regulates germline proteostasis; second, the genetic interactions of PEPT-1 with IIS and mTOR1, two significant regulators of protein synthesis, have been reported (*36–38*); finally, we showed a loss-of-function mutant, *pept-1*(lg601) was sufficient to rescue reproduction in the absence of germline HSF-1 during larval development as reduced IIS does (Fig. 6B).

Our qPCR analysis showed that the expression of *pept-1* decreased in the *daf-2(e1370)* animals and was restored in the *daf-2(e1370); daf-16(mgDf50)* double mutant (Fig. 6C), confirming that IIS activates the expression of *pept-1* by alleviating the repression by DAF-16. In contrast, *sod-3*, a Class I gene, was activated by DAF-16 when IIS was reduced (Fig. 6D). Supporting the notion that PEPT-1 functions downstream of DAF-16 in regulating germline proteostasis, inactivation of PEPT-1 partially restored fertility in the *daf-2(e1370); daf-16(mgDf50)* double mutant when HSF-1 was depleted from the germline starting at egg lay (Fig. 6E). These results strongly suggest that PEPT-1 has a vital role in the germline proteostasis pathway mediated by IIS and DAF-16.

### Insulin/IGF-1 signaling (IIS) regulates germline protein synthesis and proteostasis via peptide uptake in the intestine

PEPT-1 is located at the apical membrane of the intestine and is responsible for the uptake of di-/tri-peptides from dietary proteins (*39*). We then tested if IIS regulates germline protein synthesis via protein absorption in the intestine. Consistent with published data, RNAi against *pept-1* significantly impaired peptide uptake into the intestine (Fig. 7A&B). Similar results were observed in animals with reduced IIS. In particular, the *daf-2(e1370)* animals treated with *daf-2* RNAi significantly decreased peptide uptake (Fig. 7C&D). Furthermore, *pept-1* RNAi was sufficient to reduce the RPS-6 levels and translation rate in the germline, suggesting PEPT-1-mediated peptide uptake non-cell-autonomously promotes ribosome biogenesis and protein synthesis during gametogenesis (Fig. 7E&F, Fig. S5A&B). Finally, we tested if the change in peptide uptake impacts germline proteostasis by altering the pool of amino acids available for translation. A published study has shown that almost all free amino acids were decreased in the cytosol in the *pept-1(lg601)* animals compared to that in the wild-type (*38*). Supplement of a mixture of amino acids on the culture plates of *pept-1 (lg601)* animals has increased the number of mated progenies by 20% (P:0.06) in the presence of HSF-1 but significantly decreased the brood size upon HSF-1 depletion (Fig. 7G). Similar results were obtained when we measured self progenies (Fig. S5C). These results suggest that loss of PEPT-1 limits protein absorption and, consequently, the amino acid pool used for germline protein synthesis, which lowers the rate of translation, rendering germline development less dependent on HSF-1-mediated protein folding.

**Figure 7.**
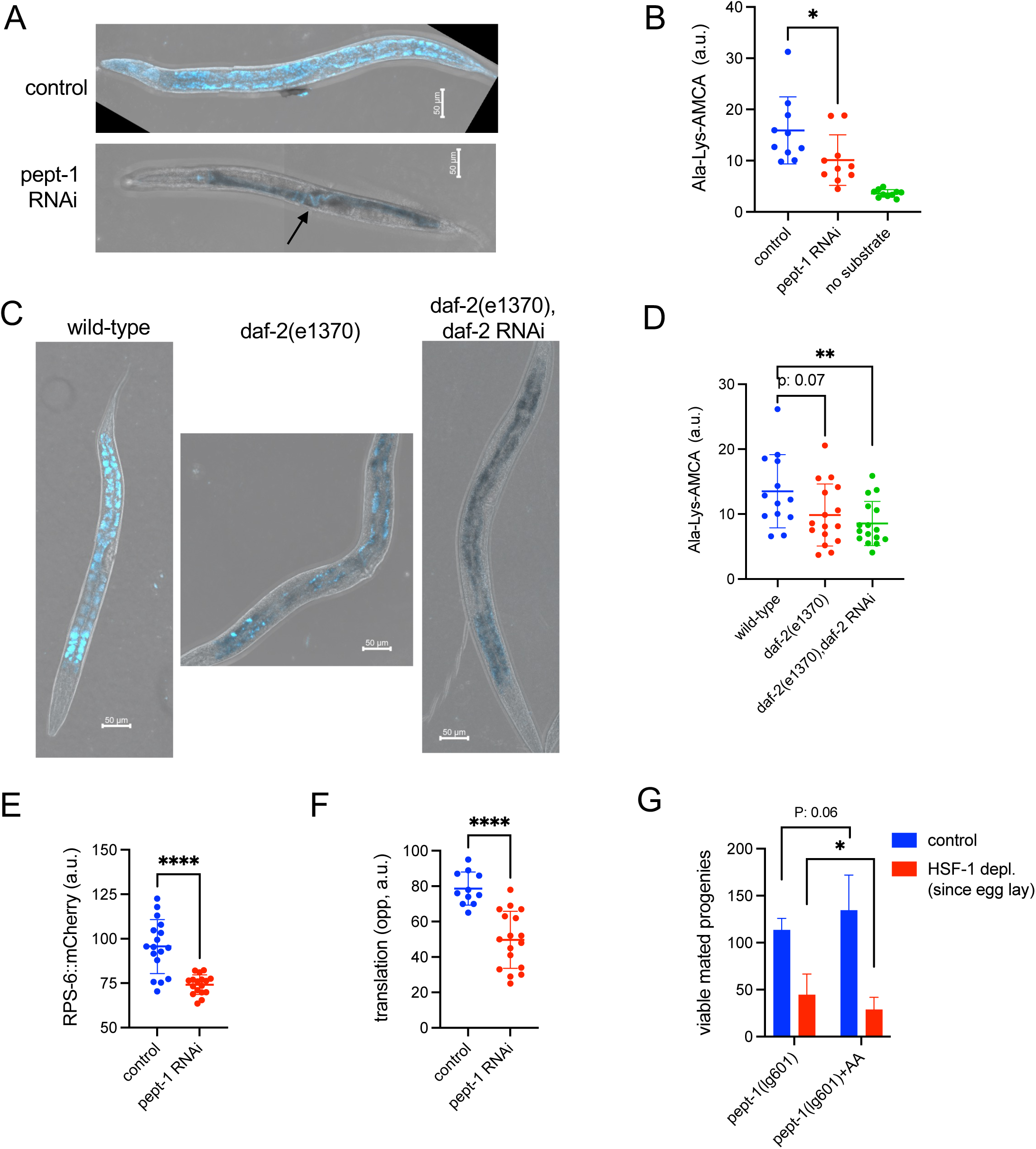
Insulin/IGF-1 signaling (IIS) regulates germline protein synthesis and proteostasis via peptide uptake in the intestine. (A&B) Representative images (A) and quantification (B) of intestinal peptide uptake measured by Ala-Lys-AMCA upon pept-1 RNAi. RNAi started at egg lay, and peptide uptake was measured in late L4 and young adult stages. The signal in the pept-1 RNAi image largely came from Ala-Lys-AMCA trapped in the lumen (as pointed by the arrow). Mean and standard deviation are plotted (N=10). Student’s t-test. *p<0.05; ns: p>=0.05. (C&D) Representative images (C) and quantification (D) of intestinal peptide uptake in animals with reduced IIS. Experiments were done as in A&B. The wild-type animals treated with control RNAi were used as the positive control. Mean and standard deviation are plotted (wild-type: N=13; *daf-2(e1370)* experiments: N=16). Student’s t-test. **p<0.01; ns: p>=0.05. (E&F) Quantification of the endogenously tagged RPS-6::mCherry protein (E) and translation measured by OPP incorporation (F) in the germline upon pept-1 RNAi. RNAi started at egg lay, and measurement was done in young adults. Mean and standard deviation are plotted (RPS-6 experiments: N=17; translation: control N=10, pept-1 RNAi N=17). Student’s t-test. ****p<0.0001. (G) Brood size analysis of the *pept-1(lg601)* animals in the presence or absence of amino acid (AA) supplements. Both depletion of HSF-1 from the germline and the supplement of AA started from egg lay. The *pept-1(lg601)* animals were mated with N2 males on Day 1 of adulthood. Mean and standard deviation are plotted (N>=12). Student’s t-test. *p<0.05.

## Discussion

Proteostasis is the cellular state in which protein synthesis, folding, transport, and turnover are well coordinated to maintain a functional and dynamic proteome (*40*). Gametogenesis involves periods of very active protein synthesis and is particularly dependent on proteostasis (*3, 41*). On the other hand, gametogenesis is highly energy-demanding and sensitive to energy metabolism and environmental stress, which impact both genome and proteome stability. Compared to the extensively studied mechanisms that safeguard germline genome integrity, how animals regulate germline proteostasis at the organismal level is still poorly understood. In this study, we took *C. elegans* as a model and demonstrated that HSF-1-dependent protein folding capacity needs to be coupled with the rate of protein synthesis controlled by insulin/IGF-1 signaling (IIS) to achieve germline proteostasis. Especially, IIS activates the expression of the intestinal peptide transporter, *pept-1,* by alleviating its repression from DAF-16, which provides a non-cell-autonomous mechanism that controls germline ribosome biogenesis and translation via regulation of dietary protein absorption (Fig. 8).

**Figure 8.**
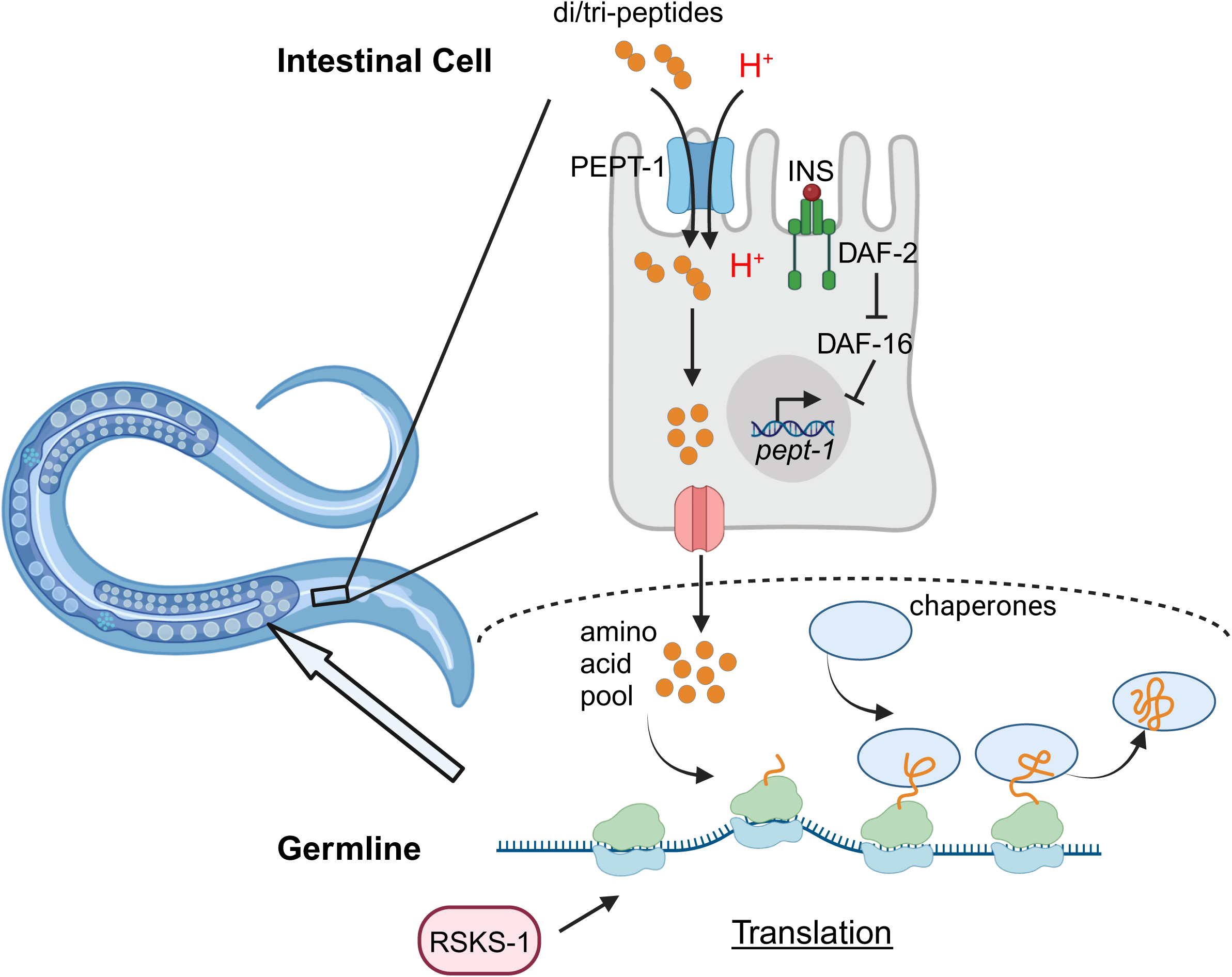
Model for non-cell-autonomous regulation of germline proteostasis by insulin/IGF-1 signaling (IIS). IGF-1R/DAF-2 activates the expression of the *pept-1* gene by alleviating its repression by FOXO/DAF-16, which promotes the uptake of di/tri-peptides from the dietary proteins. This pathway non-cell-autonomously increases the pool of amino acids that fuel ribosomal biogenesis and translation in the germline, which is the primary site of protein synthesis in reproductive animals. HSF-1-dependent chaperone expression must be at a delicate balance with IIS activity, providing sufficient folding capacity for nascent proteins. Reduced IIS lowers ribosomal biogenesis and translation, rendering the germline resilient against limited protein folding capacity and proteotoxic stress.

### HSF-1 works in concert with the insulin/IGF-1 signaling (IIS) pathway to provide sufficient protein folding capacity in germ cells

HSF-1 is best known for its ability to respond to proteotoxic stress such as heat shock and induce the expression of a series of protein quality control factors including molecular chaperones, detoxification enzymes, and players in protein clearance pathways to restore proteostasis (*42*). Published works from us and colleagues have shown that only a subset of germ cells can induce the classical heat shock response (HSR) upon temperature stress (*9, 12*). Instead, HSF-1 promotes the expression of selective chaperones and co-chaperones that are important for the folding and maturation of nascent proteins in physiological conditions (*9*). Although HSF-1 also binds and activates other genes in germ cells, our data suggest that HSF-1’s primary role in gametogenesis is to provide protein folding capacity. Loss of HSF-1 led to protein misfolding and degradation, which over time exhausted the UPS, causing accumulation of protein aggregates (Fig. 1).

Interestingly, the requirement for HSF-1 in gametogenesis is determined by the activity of IIS. Using a reduction-of-function mutant of the IGF-1R/DAF-2 and RNAi, we found the defects in fecundity, gamete quality as well as proteostasis caused by HSF-1 depletion, negatively correlated with IIS activity (Fig. 3A&B, Fig. 3F&G). As IIS increases the rate of protein translation in the germline (Fig. 4E), we propose that HSF-1-dependent chaperone expression needs to be coupled with IIS-activated protein synthesis to ensure sufficient folding capacity for nascent folding and protein maturation during gametogenesis. The mechanism of coupling likely occurs at the regulation of HSF-1 transcriptional activity. Our previous work has shown that essential chaperone genes like *hsc-70* and *hsp-90* can be expressed at the basal level without HSF-1, but HSF-1 tunes up their expression in response to IIS. This explains why only *hsp-90*, *hsc-70*, *dnj-13,* and *sti-1* significantly decreased expression upon HSF-1 depletion in the *daf-2(e1370)* mutant (Fig. 3D). These four genes showed the most prominent fold-change upon HSF-1 depletion in the wild-type animals (Fig. 1A, 16 h time point). The changes on the other chaperone genes likely fell below detection sensitivity with the lower HSF-1 activity in the germline upon reducing IIS.

However, our previous work did not answer how IIS stimulates HSF-1 activity in germ cells. Our finding in this study that IIS activates germline protein synthesis provides a plausible mechanism. It is known that chaperones, including HSC-70, HSP-90, and the TRiC/CCT complex, interact with HSF-1 and inhibit its activity (*43–45*). When IIS activity is high, the consequent high influx of nascent proteins titrates chaperones from HSF-1, releasing HSF-1 from the inhibitory state to upregulate chaperone expression and augment protein folding capacity. Coupling IIS and HSF-1 activities provides a strategy to ensure sufficient chaperone expression for nascent folding. This mechanism may have additional meanings in gametogenesis as molecular chaperones have evolved specialized roles in meiosis. For example, in *C. elegans*, both IIS and HSP-90 are required for MAPK activation during oocyte growth (*27, 46*). In future studies, it is interesting to determine how HSF-1-dependent hsp-90 expression may contribute to IIS-MAPK signaling in gametogenesis.

### The low rate of ribosome biogenesis and translation underlies robust germline proteostasis mediated by reduced insulin/IGF-1 signaling (IIS)

Animals with reduced IIS were more resilient to proteotoxic challenges associated with loss of HSF-1 from the germline (Fig. 3F&G) and by acute and chronic stresses (Fig. 5D&E). It is well-established that reduced IIS improves proteostasis and extends lifespan by enhancing stress responses in somatic tissues (*29*). This is not likely the primary mechanism underlying the robust germline proteostasis by reduced IIS. First, although the expression of HSC-70 and HSP-90 were significantly down-regulated in the *daf-2(e1370)* animals upon chronic HSF-1 depletion (Fig. 3C), it did not induce a stress-response signature based on our transcriptomic analysis (Fig. 3D&E). Second, the two stress-responsive factors downstream of IIS, NRF/SKN-1 and FOXO/DAF-16, were not required from the germline by reduced IIS for reproduction in the absence of HSF-1 (*9*). NRF/SKN-1 activates the antioxidant response (*30*), and FOXO/DAF-16 promotes the expression of chaperones, UPS components, and other protein quality control genes upon stress (*19, 28, 47, 48*). Most importantly, lowering the rate of translation in the *rsks-1(ok1255)* mutant phenocopied the robust germline proteostasis as observed in animals with reduced IIS when challenged by HSF-1 depletion or by stresses (Fig. 5). Collectively, these data provide compelling evidence that tuning down protein synthesis by reducing IIS confers resilience against limited protein folding capacity and proteotoxic stresses in gametogenesis.

Our finding is consistent with the role of IIS in nutrient and stress sensing and with the previous findings of IIS in larval development (*49*). In favorable conditions, IIS is high, speeding up development and reproduction, while under adverse conditions, IIS decreases, promoting survival and stress resilience. However, our findings suggest that the somatic tissues and the germline utilize different strategies to enhance proteostasis upon IIS reduction. The germline is the primary site of protein synthesis in reproductive animals. Therefore, tuning down ribosome biogenesis and translation could be sufficient to slow down the protein life cycle and spare chaperones and protein clearance pathways to cope with damaged and misfolded proteins upon stress. In addition, meiotic cells in the diakinesis state undergo chromosome compaction and transcription repression, limiting transcriptional stress responses. Conversely, somatic tissues may require the induction of additional protein quality control factors through transcriptional responses to cope with proteotoxic stresses. It is important to note that our RNA-seq data suggest that reduced IIS down-regulates almost all the ribosomal protein genes and multiple factors that function at translation initiation and translation elongation (Fig. 4B). Knock-down of individual translation factors such as the initiation factor, *ifg-1* was sufficient to decrease translation but not able to rescue gametogenesis upon HSF-1 depletion from the germline (data not shown). Although we cannot rule out the impacts of RNAi efficiency, this result implies that tuning down protein synthesis coordinately at ribosome biogenesis and both initiation and elongation steps of translation may be necessary for enhanced proteostasis.

Interestingly, reduced IIS is known to improve oocyte quality in maternal aging, which requires the activity of DAF-16 non-cell-autonomously from the muscle and the intestine, the same tissues where DAF-16 is required for dictating the resilience against HSF-1 depletion from the germline (Fig. 6A). As loss of proteostasis is a hallmark of aging, it raises an intriguing possibility that reduced IIS may delay oocyte aging through enhancing germline proteostasis. Future studies will determine if PEPT-1 has a role in reproductive aging and identify the signal of DAF-16 from the muscle that regulates germline proteostasis. Notably, IIS can also cell-autonomously impact germline proteostasis. A previous study shows that IIS regulates oocyte quality in aging via the Cathepsin B protease activity (*50*).

### Insulin/IGF-1 signaling (IIS) regulates germline proteostasis non-cell-autonomously via PEPT-1

Based on the following evidence, our work suggests that IIS regulates protein synthesis in the germline non-cell-autonomously via the intestinal oligo-peptide transporter, PEPT-1. First, IIS activates the expression of *pept-1* by alleviating its repression from DAF-16 (Fig. 6C). The *pept-1* promoter contains DAE (DAF-16-associated element), the proposed binding site of PQM-1, a transcription factor whose activity is antagonized by DAF-16 (*35*). The ModEncode ChIP-seq data suggest that PQM-1 binds to the *pept-1* promoter (*51*), but its role in regulating *pept-1* expression is yet to be determined. Second, the loss-of-function mutant *pept-1(lg601)* partially rescued reproduction in both the wild-type and *daf-2(e1370); daf-16(mgDf50)* double mutant when HSF-1 was absent from the germline through larval development (Fig. 6B&E) and was sufficient to reduce ribosome biogenesis and translation rate in the germline (Fig. 7E&F). It argues that PEPT-1 functions downstream from IIS and DAF-16 to regulate germline proteostasis. Finally, reduced IIS decreased the peptide uptake (Fig. 7C&D). Dietary proteins are hydrolyzed to oligopeptides in the intestinal lumen and further processed into di-/tri-peptides and free amino acids to enter enterocytes via separate transporters (*39*). *C. elegans* PEPT-1 is the ortholog of the low-affinity, high-capacity di-/tri-peptide transporter PEPT1 in mammals (*52*), which serves as one major protein absorption pathway. Supplementation of a mixture of amino acids in the culture increased the brood size of the *pept-1(lg601)* animals by ∼20% in the presence of HSF-1 (Fig. 7G&S5C), which is consistent with previous results (*36, 53*) as increasing the uptake of free amino acids through the corresponding transporters would not completely replace the function of PEPT-1. However, this was sufficient to increase the sensitivity of the *pept-1(lg601)* animals to HSF-1 depletion from the germline. It indicates that IIS through PEPT-1 promotes dietary protein absorption, which is subsequently incorporated into a pool of amino acids to fuel germline protein synthesis (Fig. 8).

It has been reported that PEPT-1 genetically interacts with IIS and mTORC1 pathways to regulate development, reproduction, and longevity (*36–38*). However, those studies have focused on the intestinal cells or the organismal functions. As the germline is the primary site of protein synthesis in reproductive animals and amino acids stimulate mTORC1 activity via multiple signaling pathways (*54*), we propose that PEPT-1-dependent amino acid sufficiency could regulate mTORC1 in the germline and impact translation. Our results on RSKS-1, the key regulator of translation downstream of mTORC1 (*55, 56*), support this hypothesis. We found that the *rsks-1(ok1255)* animals phenocopied the reproductive behaviors of animals with reduced IIS in response to HSF-1 depletion and proteotoxic stresses (Fig. 5), linking mTORC1 to IIS-regulated germline proteostasis. Future work will test if autophagy, another protein quality control pathway downstream of mTORC1 (*56*), is also involved.

PEPT-1 is a rheogenic H^+^-dependent carrier that causes an influx of proton during transport, and it requires the sodium-proton antiporter NHX-2 to recover from the intracellular acid load (*57*). NHX-2 also promotes free fatty acid uptake; thus, PEPT-1’s activity impacts fatty acid metabolism and storage (*58*). A recent study indicates that the residency of PEPT-1 transporter at the plasma membrane is regulated by lipid homeostasis in the intestine (*59*). Our work opens the opportunity for future study to understand how fatty acid and protein metabolisms interact with the glucose-sensing IIS to regulate germline proteostasis at the organismal level.

## Materials and Methods

### Experimental Models

Unless stated, *C. elegans* strains were maintained at 20°C on NGM plates seeded with OP50 bacteria and were handled using standard techniques (*60*).

The HSF-1 AID allele, *hsf-1(ljt3[hsf-1::degron::gfp])I,* was from our previous work (*9*). To perform HSF-1 depletion in strains that express NMY-2::GFP (*zuIs45 [nmy-2p::nmy-2::GFP + unc-119(+)] V*) (*61*) and GFP::H2B (*oxIs279 [pie-1p::GFP::H2B + unc-119(+)]II*) (62), we made another HSF-1 AID *allele hsf-1(ljt5[hsf-1::degron::3xFLAG])I* using CRISPR knock-in of degron::3xFLAG to the C-terminus of endogenous *hsf-1* gene through microinjection of chemically modified synthetic sgRNA (Synthego) along with Cas9 Nuclease (Integrated DNA Technologies, IDT) following the previously published protocol (*63*). The repair template was made by NEB assembly of upstream flanking sequence and degron from the repair template of degron::gfp and a synthetic gene fragment from IDT that contains a 3xFLAG tag and the downstream flanking sequence. The new HSF-1 AID model was outcrossed 6 times before use.

The Bristol N2 and other worm strains were obtained from the *Caenorhabditis elegans* Genetics Center (CGC). The alleles used in this study include:

*ieSi38[sun-1p::TIR1::mRuby::sun-1_3’UTR+Cbr-unc-119(+)]IV* (induce AID in the germline) *daf-2(e1370) III, pept-1(lg601)X, daf-16(mgDf60)I, rsks-1(ok1255)III, rde-1 (mkc36) V JAR16 rps-6(rns6[rps-6::mCherry])I (measurement of endogenous RPS-6)*

*mkcSi13 [sun-1p::rde-1::sun-1 3’UTR + unc-119(+)]II* (germline RNAi)

*neIs9 [myo-3::HA::rde-1 + rol-6(su1006)]X (*muscle RNAi)

*kzIs9 [(pKK1260) lin-26p::NLS::GFP + (pKK1253) lin-26p::rde-1 + rol-6(su1006)]* (hypodermis RNAi)

*frSi17[mlt-2p:rde-1]II* (intestine RNAi)

### HSF-1 Depletion and RNAi

HSF-1 depletion was done in the AID models that degraded the HSF-1 protein, specifically in the germline, upon auxin treatment. Auxin treatment was performed by transferring worms to bacteria-seeded NGM plates containing 1 mM auxin (indole-3-acetic acid, Sigma). The preparation of auxin stock solution (400 mM in ethanol) and auxin containing NGM plates was done as previously described (*64*). In all experiments, worms were also transferred to NGM plates containing 0.25% ethanol (EtOH) as the mock-treated control.

RNAi was performed by feeding, and all RNAi clones from the Ahringer Library (*65*) were sequence-verified before use. An RNAi-compatible OP50 strain (*66*) was used in all the experiments. Overnight cultures of RNAi bacteria in LB media containing 100 μg/ml ampicillin were diluted and allowed to grow for another 5-6 hours at 37 °C to reach OD600 of 1.0-1.2. Cultures were then seeded onto NGM plates containing 100 μg/ml ampicillin and 1 mM IPTG to induce expression of dsRNA and dry at room temperature for 2 days. All RNAi experiments were done by laying eggs directly on freshly prepared RNAi plates. Tissue-specific RNAi were performed in transgenic worms expressing the Argonaute protein gene *rde-1* in the germline or specific somatic tissue in the null mutant *rde-1 (mkc36)* (*67*).

### Translation Assay

The O-propargyl-puromycin (OPP) translation assay was performed as previously described (68), except that fixation and conjugation of fluorescent azide were done in dissected gonads rather than in the whole worms. During incubation with OPP, worms were fed with dead bacteria. OP50 bacteria were killed by incubating with 1% paraformaldehyde for 1 hour and washed three times with M9 buffer. In each experiment, 50-60 young adult worms synchronized by egg lay were used. Worms were incubated in 10µM OPP (Invitrogen) diluted with M9 buffer containing 2×10^8^ CFU/mL of dead OP50 in a total volume of 1 mL for 3 hours at 20 °C with gentle shaking. Animals were then dissected in PBS buffer containing 0.1% Tween-20 (PBST) and 0.25 mM levamisole to expose gonads. The samples were then fixed in PBST buffer containing 3% paraformaldehyde for 10 min at room temperature and then permeabilized with cold methanol for 15 min at -20°C. After three washes with PBST, the samples were used for conjugation of fluorophore to the puromycylated peptides using the Click-iT ® Plus Alexa 647 Fluor ® Picolyl Azide tool kit (Invitrogen). The pellet of gonads was resuspended and incubated with 200 µL Click-iT ® reaction buffer containing 0.5 µL of 500 µM Alexa Fluor ® Picolyl Azide, 8 µL 100mM copper protectant, 20 µL of reaction buffer additive, 17 µL of 10x reaction buffer and 153 µL of pure water for 1 hour at 20 °C with gentle shaking. After the Click-iT ® reaction, the pellet was washed with reaction rinse buffer (3% bovine serum albumin in PBST) and, subsequently, three times with PBST to remove the unconjugated Alexa Fluor ® Picolyl Azide. Finally, DNA staining was done in 100 ng/ml of DAPI for 20 min, and the samples were mounted on an agarose pad for confocal imaging.

### Proteotoxic Stress Test

The F1 survival assay following acute heat shock (HS) was performed as previously described (*69*) with some modifications. Day 1 gravid adults were exposed to a 15-minute HS at 34 °C by submerging the culture plates in a water bath and immediately moved back to 20 °C to recover. Eggs laid within 4 hours post-HS were scored for embryo lethality.

Proteotoxic stress tests by proteasome inhibitor and paromomycin were done on NGM plates seeded with paraformaldehyde-killed OP50. 400 µL M9 buffer containing 10 µM of the proteasome inhibitor, bortezomib (Sigma-Aldrich), or 1 mM of paromomycin (Sigma-Aldrich) was applied to the surface of 3.5 cm seeded NGM plates. Animals were singled to those plates at the young-adult stage and embryo lethality in the progenies was measured through the reproductive period.

### Fecundity and Embryo Lethality Measurement

As specified in figure legends, animals were synchronized by egg lay on normal NGM plates or plates containing auxin and/or RNAi bacteria. Worms were singled at the young adult stage and allowed egg-laying for 24 hours. Worms were transferred to new plates daily, and eggs were allowed to hatch and grow to the L3 stage. The number of viable larvae and dead eggs was counted at this point. In mating experiments, both males and hermaphrodites at the young-adult stage were used. The N2 males were added to hermaphrodites in a 2:1 ratio and kept for 24 hours. Only those hermaphrodites that have successful mating (close to 50% male progenies) were included in fecundity and embryo lethality analysis. Amino Acid supplementation was performed as previously described (*53*). A 1:1 mixture of MEM non-essential amino acids (100x) and MEM amino acids (50x) without L-glutamine (Sigma-Aldrich) was applied to the surface of 3.5 cm NGM plates. The freshly prepared plates were used the next day.

### Germline Nuclei Isolation

Germline nuclei isolation was performed by following a published protocol (*70*) except that worms were lightly fixed in 50 mL of −20°C dimethylformamide (Sigma-Aldrich) for 2 min and washed three times in PBS before homogenization. This light fixation helped preserve the RNA from degradation. About 25,000 young adult worms from fifty 10cm NGM plates were used in each nuclei prep. Both the enriched germline nuclei and whole worm lysate were saved for RNA-seq analysis.

### RNA Extraction, cDNA Synthesis, and qPCR

Animals were synchronized by treatment of alkaline hypochlorite solution or egg laying (for auxin treatment starting from egg lay). When using alkaline hypochlorite, synchronized L1 larvae were grown on 10 cm normal NGM plates (∼500 worms per plate) for larval development. When checking DAF-16-dependent *pept-1* expression (Fig. 6), worms were collected at the late L4 stage when *pept-1* expression peaks. For RNA-seq analysis of the *daf-2(e1370)* animals, approximately 120 adult worms were used in each experiment. The worms were either kept on 10 cm NGM plates containing either EtOH or auxin since egg lay or picked onto those plates at the young adult stage and kept for 16 hours. RNA was extracted using 300 μL Trizol reagent. Worms were vortexed continuously for 20 minutes at 4°C and then went through one cycle of freeze-thaw to help release RNA. Following this, RNA was purified using the Direct-zol RNA MiniPrep kit (Zymo Research) per the manufacturer’s instructions using on-column DNase I digestion to remove genomic DNA. RNA was used in library preparation for sequencing or to synthesize cDNA for qPCR analysis.

cDNA was synthesized using the BioRad iScript cDNA synthesis kit per the manufacturer’s instructions. Relative mRNA levels were then determined by real-time quantitative PCR using iTaq Universal SYBR Green Supermix (BioRad) and a ThermoFisher QuantStudio 6 Pro thermocycler. Relative mRNA levels were calculated using the standard curve method, and gene expression was normalized to the mean of the housekeeping genes *cdc-42* and *rpb-2*. The following primers were used in qPCR:

pept-1-F: ACTATGGAATGAGAACGGT; pept-1-R: CTTGTCCGATTGCGTAT

sod-3-F: CACTGCTTCAAAGCTTGTTCA; sod-3-R: ATGGGAGATCTGGGAGAGTG rpb-2-F: AACTGGTATTGTGGATCAGGTG; rpb-2-R: TTTGACCGTGTCGAGATGC

cdc-42-F: TGTCGGTAAAACTTGTCTCCTG; cdc-42-R: ATCCTAATGTGTATGGCTCGC

### RNA-seq Analysis

Total RNAs were polyA enriched, and directional RNA-seq libraries were prepared using the NEBNext Ultra II RNA library prep Kit. Paired-end sequencing was done at a NovaSeq 6000 sequencer at the OMRF clinical genomics core.

RNA-seq reads were mapped to Ensembl WBcel235 genome using RNA STAR (*71*) with -- alignIntronMax 120000 to set the intron size, and --outFilterMultimapNmax 200 to allow multi-mapped reads. The mapped reads were then subject to FeatureCounts in Rsubread (*72*) for quantification with the setting -p -B -P -C -M -O --fraction –largestOverlap. The settings in STAR and FeatureCounts enabled proper quantification of those heat shock genes (e.g., *hsp-70* and *hsp-16*s) duplicated in the *C. elegans* genome. Differential expression (DE) analyses were done using edgeR (*73*) with default settings except for using the Likelihood Ratio Test and filtering out those lowly expressed genes with CPM (counts per million) values less than 1 in more than 75% of samples. We then filtered out DE genes caused by auxin treatment using a list from our previous study (*9*) and reported only DE genes caused by HSF-1 depletion.

Gene ontology analysis (GO) was done using the program DAVID (http://david.abcc.ncifcrf.gov/) with functional annotation clustering to collapse redundant GO terms. The enrichment score for each cluster was shown with the corresponding GO_BP (Biological Processes) term representing the cluster.

### Peptide uptake assay

Peptide uptake was done as previously described (*36*). About 30-40 worms at the Late L4/young adult stage were picked into cold M9 buffer and washed twice to remove bacteria. Worm pellets were resuspended in M9 buffer containing 1 mM β-Ala-Lys-AMCA (peptide institute), transferred into a 96-well plate and incubated at 20°C for 3 hours with gentle shaking. Worms were washed in M9 buffer and mounted onto an agarose pad with levamisole for imaging.

### Immunofluorescence (IF)

IF was performed with dissected gonads as previously described (*74*) with slight modifications. Animals were synchronized by egg lay and grown to the young adult stage. Approximately 30 animals were dissected in PBS buffer containing 0.1% Tween-20 (PBST) and 0.25 mM levamisole. The samples were fixed in a PBST buffer containing 3% paraformaldehyde for 10 minutes at room temperature and then incubated with cold methanol for 15 minutes at -20 °C. After washes with PBST, the samples were blocked for 30 min at room temperature in PBST with 1% bovine serum albumin before the antibodies against histone H3 at 1:800 dilution (rabbit, Abcam) and ubiquitinylated proteins (clone FK2, mouse, Sigma-Aldrich) at 1:200 dilution were added. After overnight incubation at 4 °C and four washes with the blocking solution, Goat anti-Rabbit IgG (H+L) Alexa Fluor Plus 488 (Invitrogen) and anti-mouse-Alexa 647 (Invitrogen) were added to the samples at 1:1000 dilution, and the samples were incubated for 2 hours in the dark at room temperature. DNA staining was done in 100 ng/ml of DAPI for 20 min, and the samples were mounted on a 2% agarose pad for confocal imaging.

### EdU labeling

EdU labeling was performed by feeding animals with EdU-labeled MG1693 bacteria and by click reaction as previously described (*74*). Briefly, the animals were transferred onto M9 plates with EdU bacteria and fed for half an hour. Animals were dissected in PBS buffer containing 0.1% Tween-20 (PBST) and 0.25 mM levamisole, fixed for 10 minutes in 3% paraformaldehyde (Fisher Scientific) in PBST, washed in PBST, and incubated for one hour in -20°C methanol. Rehydration was done by washing the samples three times in PBST. EdU click reaction using Click-iT™ EdU Cell Proliferation Kit with Alexa Fluor™ 594 dye (Thermofisher) and DAPI staining were performed as described (*74*). Worms were then mounted on a 2% agarose pad for confocal imaging. Nuclei with any EdU labeling (individual or all chromosomes) were scored positive. Mitotic and transition zones were defined based on the crescent-shaped DAPI staining in the transition zone.

### Fluorescence Imaging and Quantification of Fluorescence Intensity

Imaging of live animals, as in Figures 1B, S1A-D, 4C, S5A, and 7A&C, was done by immobilizing animals in a drop of M9 buffer containing 6 mM levamisole on a 2% agarose pad. Images were acquired immediately using a Zeiss LSM880 or LSM980 Confocal Microscope with a 20X or 40X objective. As specified in the figure legend, proteasome inhibitor was applied to a subset of live-imaging experiments. In those experiments, worms were transferred to the M9 buffer containing 10 µM of bortezomib and paraformaldehyde-killed OP50, and incubated in 96-well plates for 6 hours at 20 °C with gentle shaking before imaging. Fluorescent imaging for IF, translation assay, and EdU labeling was performed using a Zeiss LSM880 or LSM980 Confocal Microscope through a 63X oil objective. Zen software was used to obtain z-stacks and subsequent analyses. Fluorescence intensity was quantified in individual worms after maximal intensity projection. Regions of interest were outlined within individual worms, and the arithmetic mean of fluorescence intensity per area was determined.

### Biochemical Analysis of Protein Aggregates

The fractionation and analysis of protein aggregates were performed as previously described (*75*). About 3000 age-synchronized adult worms, either with HSF-1 depletion for 24 hours or mock-treated, were harvested by washing off the plates using cold M9 buffer and washed twice to remove bacteria. The worms were washed once in ice-cold lysis buffer (20mM potassium-phosphate, pH 6.8, 1 mM DTT, 1mM EDTA, 0.1% (v/v) Tween 20 and protease inhibitor cocktail (Roche)) and frozen in liquid nitrogen. The frozen animals were ground with a plastic pestle on dry ice and then resuspended in 1 mL of ice-cold lysis buffer. The worms were further broken up using a glass Dounce tissue grinder. After successful lysis, the lysate was centrifuged for 5 min at 200 g at 4 °C to remove debris. The supernatant of multiple samples was adjusted to identical protein concentrations using BCA assay (ThermoFisher). Aggregated proteins were isolated by spinning at 18,000 g for 20 min at 4 °C. The resulting pellet was then sonicated in a Biorupter (Diagenode) using a high energy setting for 6 min with 30s on/30s off in washing buffer (2% NP-40, 20mM potassium-phosphate, pH 6.8, and protease inhibitor cocktail), and subsequently centrifuged at 18,000 g for 20 min at 4 °C. This washing step was performed twice. Aggregated proteins were then solubilized in 2% SDS sample buffer at 95 °C for 10 min. Proteins were separated by SDS–PAGE and analyzed by immunoblotting. In western blot analysis, primary antibodies against α-tubulin at 1:5000 dilution (mouse, Sigma-Aldrich) and ubiquitin at 1:2000 dilution (mouse, Novus) were used. The blot was then probed by Goat anti-Mouse Poly-HRP secondary antibody at 1:5000 dilution.

### Statistical Analysis

Two-tailed, unpaired Student’s t-tests were used for fecundity and embryo lethality comparison, RT-qPCR, and image quantifications. Error bars represent SEM or standard deviation as specified in figure legends. Benjamini and Hochberg FDR was used to calculate the adjusted P-value for differential expression analyses by RNA-seq.

## Acknowledgments

We thank Dr. Jarod Rollins for sharing worm strains, Dr. Shuai Gao for sharing equipment, and Dr. Patricija van Oosten-Hawle for helpful discussion on PQM-1. We thank members of Li Lab for providing feedback on the manuscript.

## Author contributions

Conceptualization: TM, JL; Methodology: TM, SLE, ACM, MVJ, VDO, TC, JL; Investigation: TM, SLE, ACM, MVJ, VDO, TC, SL, NGTN, JL; Data Analysis: TM, JL; Visualization: JL; Supervision: JL; Writing—original draft: JL; Writing—review & editing: TM, MJ, JL

## Data and materials availability

The RNA-seq datasets from this study have been deposited at Gene Expression Omnibus (GSE256186) and are publicly available as of the date of publication. The RNA-seq analysis on HSF-1 depletion from the germline of wild-type animals used our published data from GSE162066. Worm strains generated in this study will be deposited to the Caenorhabditis elegans Genetics Center at the University of Minnesota upon publication. Further information and requests for resources and reagents should be directed to and will be fulfilled by the Lead Contact, Jian Li.

## Supplemental files

**Figure S1, related to Figure 1.**
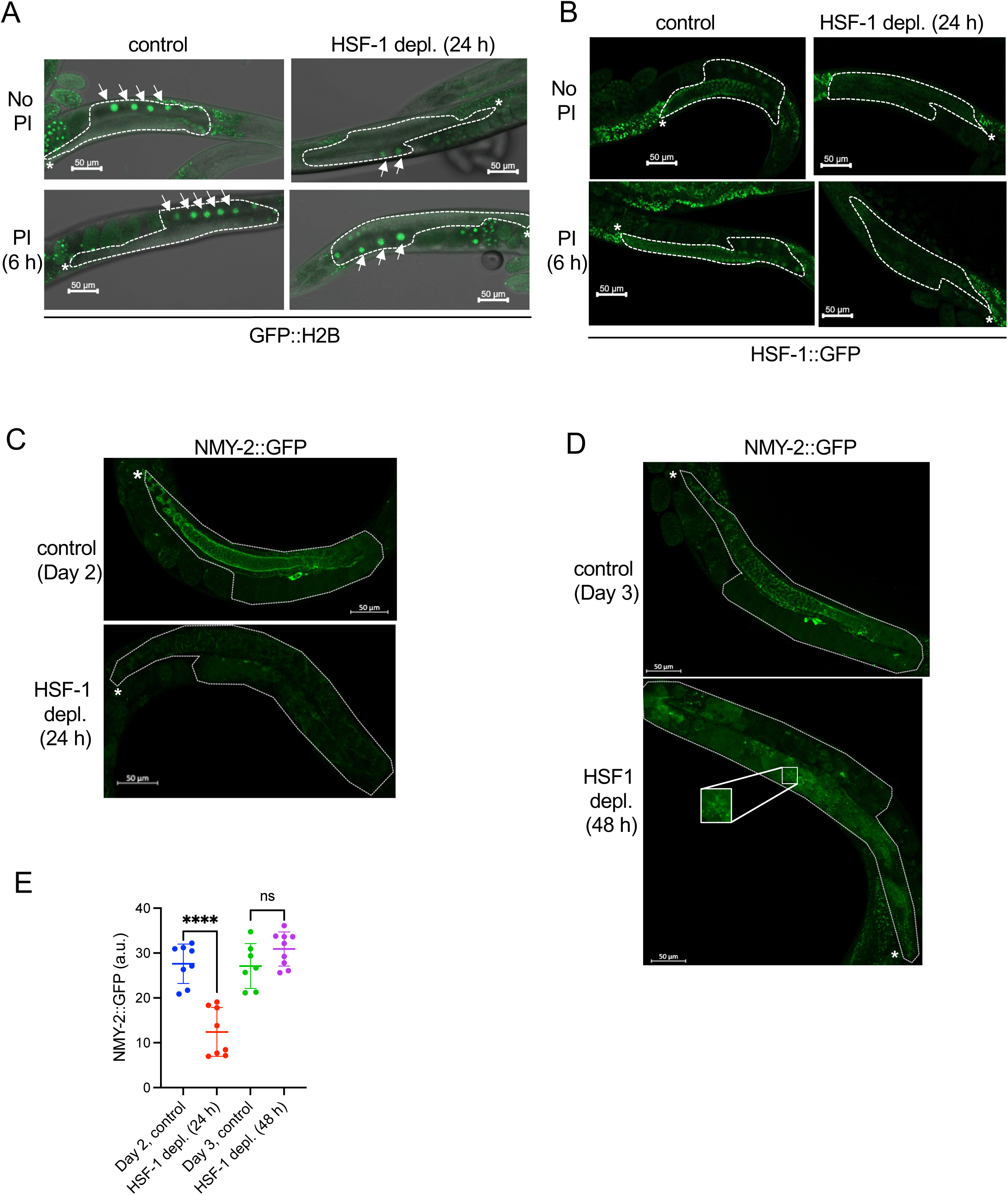
Loss of HSF-1 from the germline impairs the expression of chaperone genes, causing protein degradation and aggregation. (A) Representative images of GFP::H2B transgene in the germline upon HSF-1 depletion. HSF-1 was depleted from the germline of young adults using AID for 24 h in the presence or absence of the proteasome inhibitor bortezomib (PI) for the last 6 h of HSF-1 depletion. The dashed lines outline the gonads. The white asterisks (*) indicate the distal end of gonads where progenitor cells are located. The arrows indicate the fully grown oocytes, where the levels of GFP::H2B are quantified. (B) Representative images of the endogenously tagged HSF-1::degron::GFP upon auxin-induced degradation. HSF-1 was depleted from the germline of young adults upon auxin treatment for 24 h in the presence or absence of the proteasome inhibitor bortezomib (PI) for the last 6 h of HSF-1 depletion. The dashed lines outline the gonads. The white asterisks (*) indicate the distal end of gonads where progenitor cells are located. The proteasome inhibitor treatment was not sufficient to restore HSF-1 protein levels. (C-E) Representative images of NMY-2::GFP transgene in the germline upon HSF-1 depletion from germ cells starting at the young-adult stage for 24 h (C) and 48 h (D), and the corresponding quantification (E). The dashed lines outline the gonads. The white asterisks (*) indicate the distal end of gonads where progenitor cells are located. In the dot plots, the mean and standard deviation are plotted. Student’s t-test. ****p<0.0001; ns: p>=0.05.

**Figure S2, related to Figure 3.**
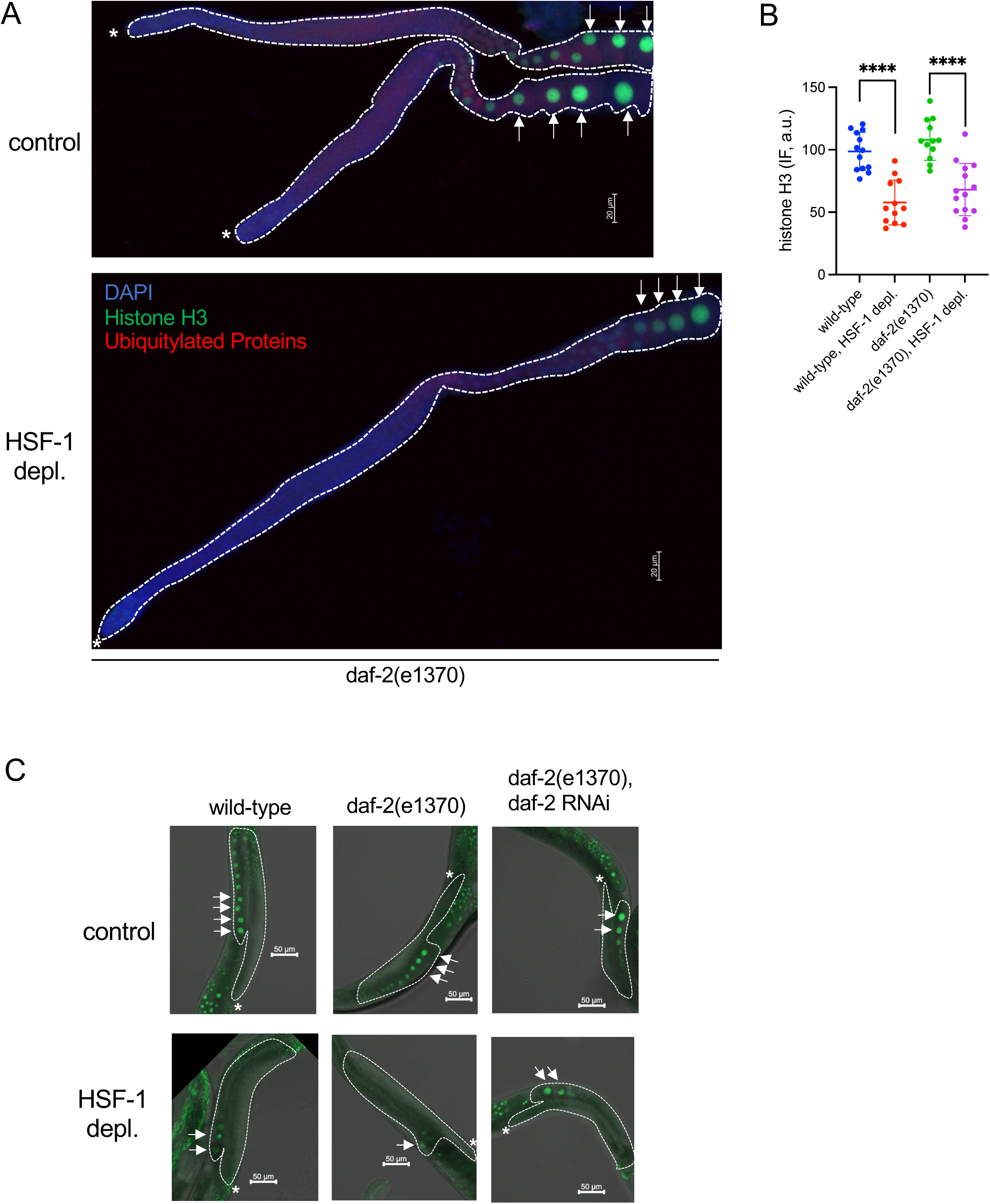
Reduced Insulin/IGF-1 signaling (IIS) confers resilience against limited protein folding capacity in gametogenesis. (A&B) Representative images (A) of endogenous histone H3 and ubiquitylated proteins (not free ubiquitin) by immunofluorescence (IF) upon germline-specific depletion of HSF-1 in the *daf-2(e1370)* animals starting from the young-adult stage for 24 h. The dashed lines outline the gonads with the white asterisks (*) marking the distal end. The levels of histone H3 in the fully grown oocytes (as indicated by arrows) are quantified (B) together with the results from the wild-type animals as a control. The mean and standard deviation are plotted. Student’s t-test. ****p<0.0001. (C) Representative images of GFP::H2B transgene in fully grown oocytes upon germline-specific depletion of HSF-1 for 24 h from young adults. The wild-type and *daf-2(e1370)* animals were treated with daf-2 RNAi or control RNAi (L4440) starting from egg lay. The dashed lines outline the gonads with the white asterisks (*) marking the distal end. The arrows indicate the fully grown oocytes, where the levels of GFP::H2B are quantified.

**Figure S3, related to Figure 4.**
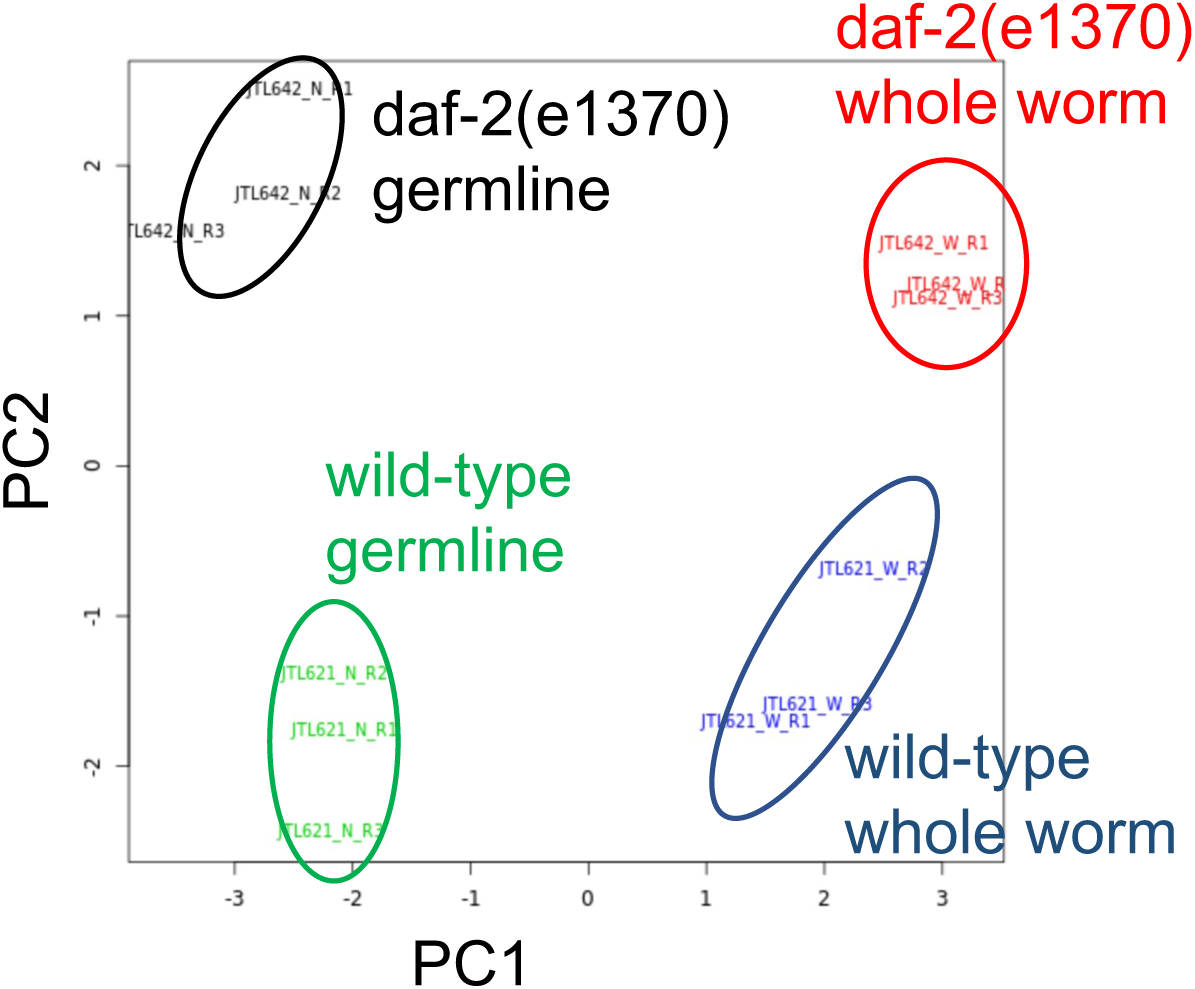
Insulin/IGF-1 signaling (IIS) activates ribosome biogenesis and translation in germ cells. Principal component analysis (PCA) on the RNA-seq results of the whole worm lysate or isolated germline nuclei from young adults of the wild-type and *daf-2(e1370)* animals (N=3).

**Figure S4, related to Figure 5.**
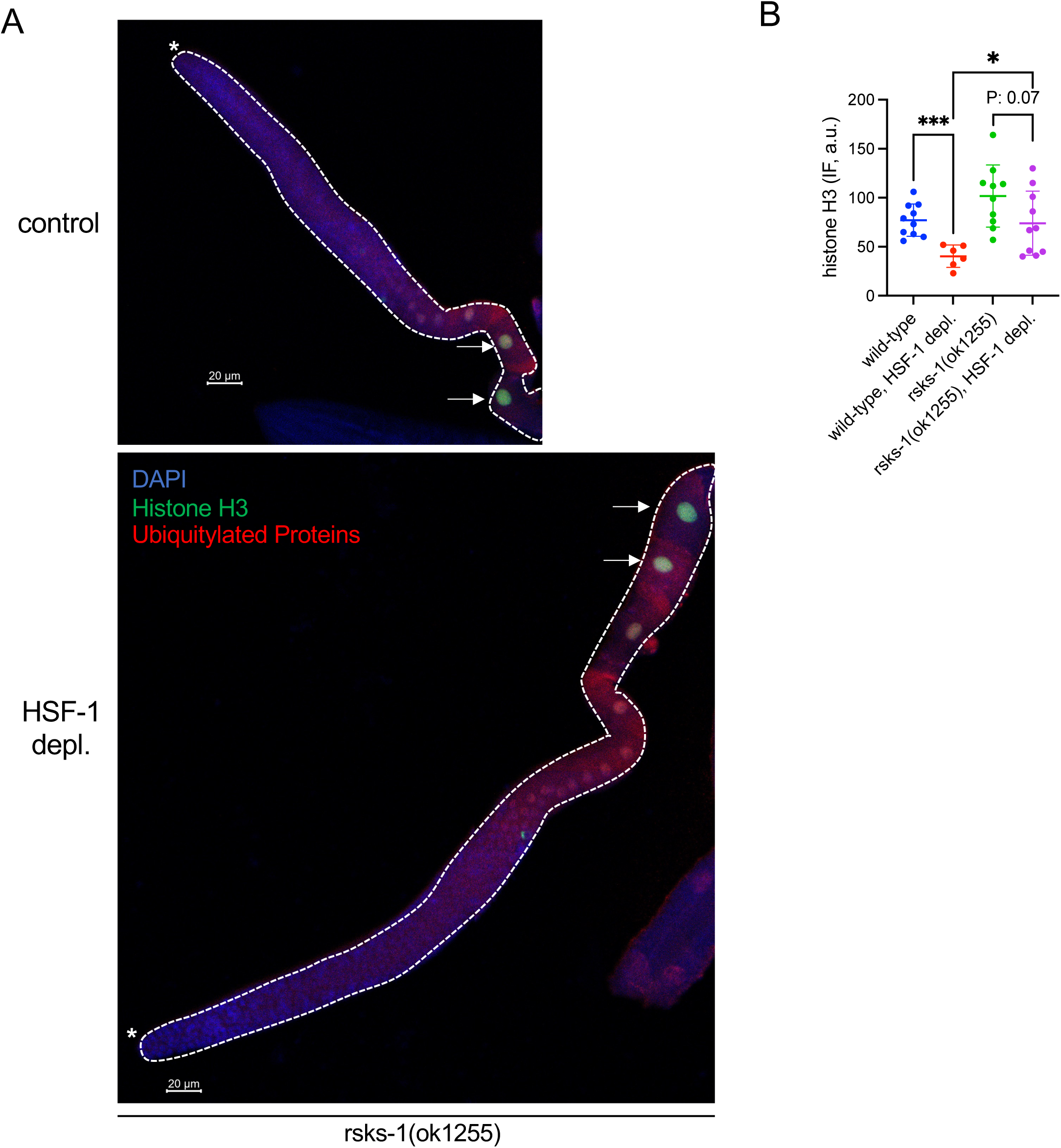
Reduced translation rate underlies robust germline proteostasis and resilience against proteotoxic stress mediated by low insulin/IGF-1 signaling (IIS). Representative images (A) of endogenous histone H3 and ubiquitylated proteins (not free ubiquitin) by immunofluorescence (IF) upon germline-specific depletion of HSF-1 in the *rsks-1(ok1255)* animals starting from the young-adult stage for 24 h. The dashed lines outline the gonads with the white asterisks (*) marking the distal end. The levels of histone H3 in the fully grown oocytes (as indicated by arrows) are quantified (B) together with the results from the wild-type animals as a control. The mean and standard deviation are plotted. Student’s t-test. ***p<0.001, *p<0.05.

**Figure S5, related to Figure 7.**
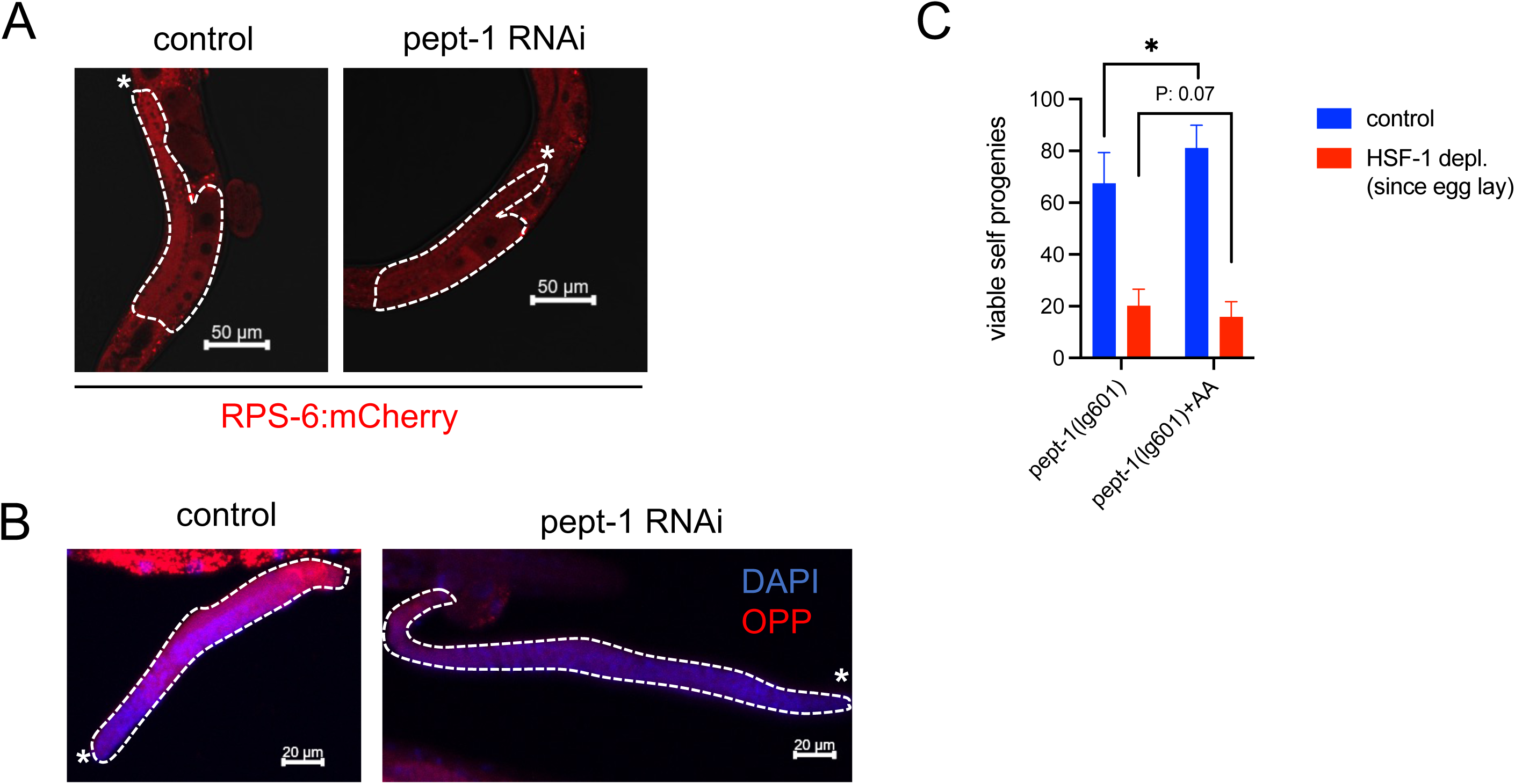
Insulin/IGF-1 signaling (IIS) regulates germline protein synthesis and proteostasis via peptide uptake in the intestine. (A&B) Representative images of the endogenously tagged RPS-6::mCherry protein (A) and translation measured by OPP incorporation (B) in the germline upon pept-1 RNAi. RNAi started at egg lay, and measurement was done in young adults. (C) Brood size analysis of the *pept-1(lg601)* animals in the presence or absence of amino acid (AA) supplements. Both depletion of HSF-1 from the germline and the supplement of AA started from egg lay. Mean and standard deviation are plotted (N>=12). Student’s t-test. *p<0.05.

